# Sequence-to-function modeling uncovers the context-specific grammar of *Drosophila* chromatin insulation

**DOI:** 10.64898/2026.06.15.732488

**Authors:** Brendan Wang, Gabriel Dolsten, Wenfan Ke, Wuwei Zhang, Anton V. Persikov, Xin Yang Bing, Xiao Li, Miki Fujioka, James B. Jaynes, Amina Kurbidaeva, Tiffany Park, Mona Singh, Michael S. Levine, Paul Schedl, Yuri Pritykin

**Affiliations:** Lewis-Sigler Institute for Integrative Genomics, Princeton University, Princeton, NJ 08544, USA; Department of Computer Science, Princeton University, Princeton, NJ 08540, USA; Center for Computational Biology, Flatiron Institute, Simons Foundation, New York, NY 10010, USA; Department of Molecular Biology, Princeton University, Princeton, NJ 08544, USA; Department of Biochemistry and Molecular Biology, Thomas Jefferson University, Philadelphia, PA 19107, USA; Center for Genomics and Systems Biology, New York University, New York, NY 10003, USA

## Abstract

Chromatin is organized into self-interacting topologically associating domains partitioned by boundary elements that insulate adjacent domains and restrict regulatory interactions. Yet, how sequence context and combinations of factors dictate boundary strength remains incompletely understood. Here we present Domino, a deep learning framework that maps genomic sequences to quantitative insulation scores defined directly from single-nucleosome resolution *Drosophila melanogaster* Micro-C data. Unlike traditional transcription factor motif scanning, Domino captures broad sequence context to resolve the functional contributions of individual sequence elements. We validate model predictions through experimental perturbations of insulator sequences. Model interpretation yields insulation-associated motifs genome-wide. Across 7,311 embryonic boundaries, Domino reveals a comprehensive insulation grammar defined by just 24 primary motifs that account for 59% of the boundaries, with an average of only two motifs per motif-containing boundary. Beyond known factors, we identify the zinc-finger proteins Trem, CG4854 and CG17385 as previously unreported insulation factors. We uncover distance– and orientation-dependent motif synergy, including a strict orientation preference of the prominent architectural factor M1BP. Finally, Domino traces tissue-specific shifts in the insulator landscape from the embryo to larval and adult brains, nominating new brain-specific insulation motifs. In sum, Domino provides a generalizable framework for decoding the regulatory logic of 3D genome architecture.

## Introduction

The genome is partitioned into self-insulating topologically associating domains (TADs), which restrict regulatory interactions between enhancers and their target genes^1–4^. This compartmentalization is enforced by sequence-specific architectural proteins that localize to TAD boundaries. While mammalian boundaries are frequently characterized by a strong reliance on CTCF and cohesin^5–7^, *Drosophila melanogaster* domain organization features a highly diversified network of distinct structural proteins operating without a single prevailing factor. In *Drosophila*, the relative contributions of transcription, chromatin accessibility, and diverse architectural proteins to insulation at boundaries remain a subject of debate^8–12^. These boundaries harbor an array of binding sites defined by sequence motifs for transcription factors (TFs) and insulator proteins, many of which remain uncharacterized^13,14^. Together, these elements establish a sophisticated insulation grammar defined by motif orientation, spacing, and combinatorial logic, that is difficult to dissect using conventional approaches^15^. Critically, since loop and distal metaloop anchors, including promoters, enhancers, and tethering elements, frequently drive or coincide with insulation, decoding this sequence grammar at high resolution, including within weak boundaries, serves as a comprehensive proxy for understanding the broader and more complex mechanisms of chromosomal organization and gene regulation^10,16–20^.

Deep learning has emerged as a powerful tool for predicting functional readouts from DNA sequence, including TF binding, chromatin accessibility, and chromatin conformation^21–25^. However, existing models for *Drosophila* chromatin architecture face significant hurdles: the compact genome provides limited training data, and prior efforts have focused on binary TAD boundary classification rather than modeling continuous levels of insulation or sequence interdependencies^14,26,27^. Recent advances in chromosome conformation capture technologies, most notably the development of Micro-C, now provide single-nucleosome resolution of chromatin features. This allows quantitative levels of insulation to be defined at high resolution directly from physical contact data, offering an unprecedented opportunity to train sequence-to-function models that can decode the precise regulatory code of insulation^16, 18, 20, 28–30^.

Here we present Domino, a deep learning framework that predicts continuous quantitative chromatin insulation directly from DNA sequence. Trained on high-resolution *Drosophila* embryo Micro-C data, Domino accurately predicts insulation levels on held-out data and across *in vivo* sequence perturbations. Model interpretation identifies motifs of both known and novel insulation factors, including the zinc-finger proteins Trem, CG4854, and CG17385, which we confirmed via experimental validation. Through genome-wide analysis, we uncover a comprehensive regulatory grammar, revealing that nearly all motif-dependent insulation is governed by just 24 primary motifs. Furthermore, *in silico* experiments reveal orientation– and distance-dependent synergies between these motifs. Finally, by applying Domino to larval and adult brain data, we trace the shifting landscape of active insulators, uncovering an increasingly prominent role for the GAGA motif and neural-specific factors in the brain. Together, these results establish Domino as a generalizable framework for deciphering the 3D regulatory logic encoded in the genome.

## Results

### Domino accurately predicts chromatin insulation directly from DNA sequence

To decode the regulatory logic of chromatin architecture, we developed Domino, a deep learning framework that predicts local insulation directly from DNA sequence (**Fig. 1A,B, Methods**). To empower this model with high-resolution experimental data, we compiled 11 Micro-C datasets from *Drosophila melanogaster* embryos at nuclear cycle 14 (nc14) (**Table S1**), yielding an aggregate map at 200 bp resolution with 1.9 billion contacts. From this map, we identified 7,311 high-confidence boundaries across the genome to serve as primary targets for analysis. Specifically, a local insulation score was computed for each 200 bp bin by using a sliding window along the contact matrix diagonal to aggregate the relative frequency of chromatin interactions crossing over a given locus, with local score minima defining the precise coordinates of the domain boundaries. To account for differences in absolute insulation across genomic regions, we applied a mean-scaling procedure to output a normalized insulation score, hereafter referred to as “insulation score” or “insulation.”

**Figure 1.**
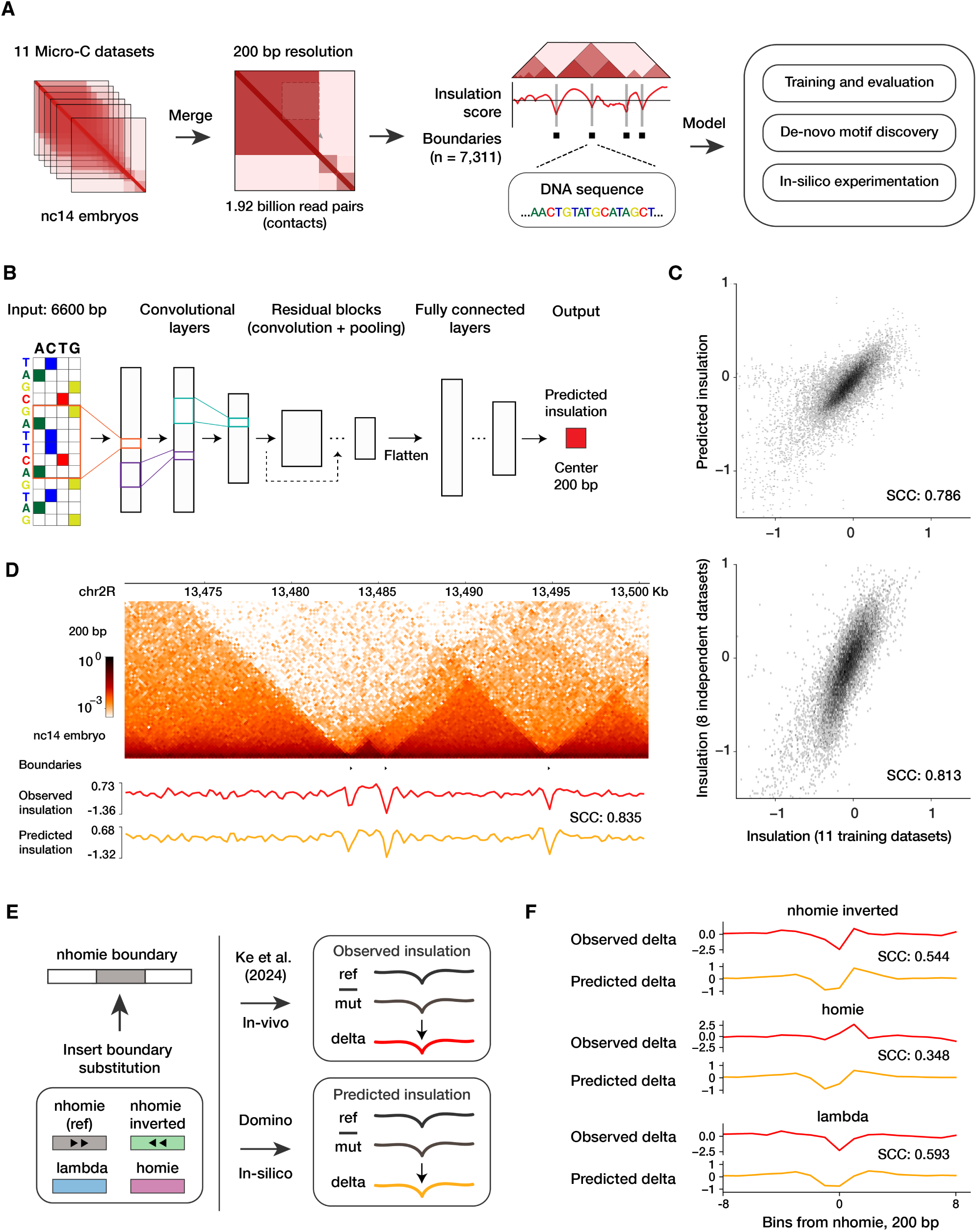
Domino: a high-resolution deep learning framework for genome-wide chromatin insulation prediction. (**A**) Overview of the Domino framework. Chromatin insulation score was defined at 200 bp resolution using 11 *Drosophila melanogaster* nc14 embryo datasets (1.9 billion total read pairs), yielding 7,311 boundaries (more negative scores indicate stronger insulation). Domino, a deep learning model, was trained on a balanced set of these boundaries and non-boundary control regions, then tested on held-out data. The trained model enables quantitative insulation prediction for arbitrary DNA sequences, facilitating *de novo* insulation motif discovery and *in silico* dissection of synergistic motif contributions to chromatin insulation (**Methods**). (**B**) Model architecture. Domino is trained to predict a normalized insulation score (200 bp bin) at the center of a 6.6 Kb input DNA sequence. (**C**) Model performance on held-out test data. Scatterplots show comparison between predicted and observed insulation (top) and between biological replicates (bottom) across held-out examples. The replicate comparison (bottom) was performed between the 11 training datasets and 8 independent Micro-C datasets not used in Domino training. SCC, Spearman correlation coefficient. For visualization, outliers representing 0.27% (top) and 1.16% (bottom) of data points are not shown. (**D**) A representative track showing observed and predicted insulation scores across a 30.2 Kb held-out genomic region. (**E**) Validation of Domino via *in vivo* sequence perturbations at the *nhomie* boundary using experimental data from Ke *et al.* 2024^30^. The central 600 bp of *nhomie* was perturbed through inversion, substitution with a control *lambda* fragment, or substitution with the *homie* boundary sequence. Insulation changes (delta) following perturbation in each mutant strain compared with the reference control were calculated from experimental Micro-C data and compared against Domino *in silico* predictions. Arrows indicate the orientation of the *nhomie* sequence. (**F**) Comparison between predicted and observed insulation changes in the three *nhomie* mutants described in (E). The 1.8 Kb region centered on the mutation site is shown.

Domino uses a combination of convolutional layers, residual blocks, and fully connected layers to map a 6.6 Kb DNA input to a quantitative insulation score for the central 200 bp bin (**Fig. 1A,B**). We performed an exhaustive hyperparameter sweep to identify the optimal architecture (**Fig. S1A, Methods**). Domino was trained on a balanced set of sites with a broad range of insulation scores and sites without insulation (**Methods**). On a held-out test set, predictions achieved a Spearman correlation of 0.79 (**Fig. 1C**). To benchmark this performance against the inherent noise in the experimental data, we analyzed 8 additional independent nc14 Micro-C datasets that were withheld from all stages of model training and selection. Notably, the correlation between Domino predictions and the experimental ground truth was on par with the correlation observed between these 8 independent replicates and the 11 training datasets (**Fig. 1C, Methods**). This suggests that Domino has captured the sequence-to-insulation relationship to a degree that approaches the theoretical limit defined by experimental variability. To evaluate the model’s ability to reconstruct the insulation landscape across continuous genomic intervals, we generated predictions for consecutive 200 bp bins genome-wide. This approach accurately recapitulated the experimental insulation tracks in held-out regions, achieving median correlations of 0.81, on par with biological replicates (**Fig. 1D, Fig. S1B**), confirming that Domino successfully generalized the sequence determinants of insulation genome-wide. Notably, correlations of insulation tracks between biological replicates exceeded those of the underlying 2D Micro-C contact maps, suggesting that sliding window-based data aggregation used in insulation analysis helps mitigate experimental noise inherent in Micro-C data (**Fig. S1B**).

To determine if Domino could generalize to sequences not present in the native genome, we validated the model using published data from *in vivo* sequence perturbations of the canonical *nhomie* boundary^30^. This experimental dataset included Micro-C for four distinct 600 bp modifications: a reference re-insertion of the original *nhomie* sequence, an inversion, a substitution with the *homie* boundary, and a substitution with a neutral *lambda* phage sequence (**Fig. 1E**). Although these perturbations were profiled in stage 14 (st14) embryos, the contact map and insulation profiles at this locus remained consistent with the nc14 data used for model training, justifying the use of st14 data for this validation (**Fig. S1C,D**). Remarkably, insulation changes derived from Domino predictions were consistent with the experimental ground truth across all mutant strains (**Fig. 1F**). This analysis provides experimental validation of Domino accuracy and its capacity to generalize beyond the endogenous sequences.

These results establish Domino as a robust framework for modeling chromatin insulation, demonstrating high accuracy, generalizability to unseen sequences, and strong agreement with experimental *in vivo* perturbations.

### Interpretation of Domino reveals the motif grammar of chromatin insulation

Having established Domino as a robust predictive model, we used it to identify the sequence features of insulatory activity. We hypothesized that such features would correspond to the binding motifs of architectural factors.

We first computed base-pair resolution attribution scores for all 7,311 boundaries using *in silico* mutagenesis (ISM)^32,33^ (**Fig. 2A, Methods**). Attribution signal was predominantly concentrated within the central 400 bp of the boundary positions (**Fig. 2B**). At high resolution, these attribution tracks frequently localized to known insulatory motifs, such as M1BP and Motif-6 (**Fig. 2A**)^14,34,35^. To systematically decode these patterns, we applied the TF-MoDISco algorithm^31^ to cluster high-importance sequences (“seqlets”) within 600 bp of the identified boundaries into 24 distinct motifs with negative attributions whose presence is predicted to strengthen insulation, identifying these motifs as the primary predicted drivers of insulation (**Fig. 2, Methods**). (Interestingly, two of these motifs also had positive attributions in some boundaries, where their presence weakens insulation (**Fig. S2**).) The majority of the motifs were associated with known architectural factors. Many of them corresponded to factors with established insulatory activity such as Su(Hw), CTCF, and the CP190-recruiting factors M1BP, Pita, Ibf, and BEAF-32^13,34,36–41^. Notably, Domino also identified motifs of recently described distal tether-element looping factors Vostok and Mulberry^42,43^, and multi-functional factors Zelda and GAF with particular activity during early development^44,45^. These attributions highlight an additional role for these looping and developmental factors as contributors to local chromatin insulation, reinforcing the idea that insulation frequently coincides with other features of 3D chromatin architecture and transcriptional regulation.

**Figure 2.**
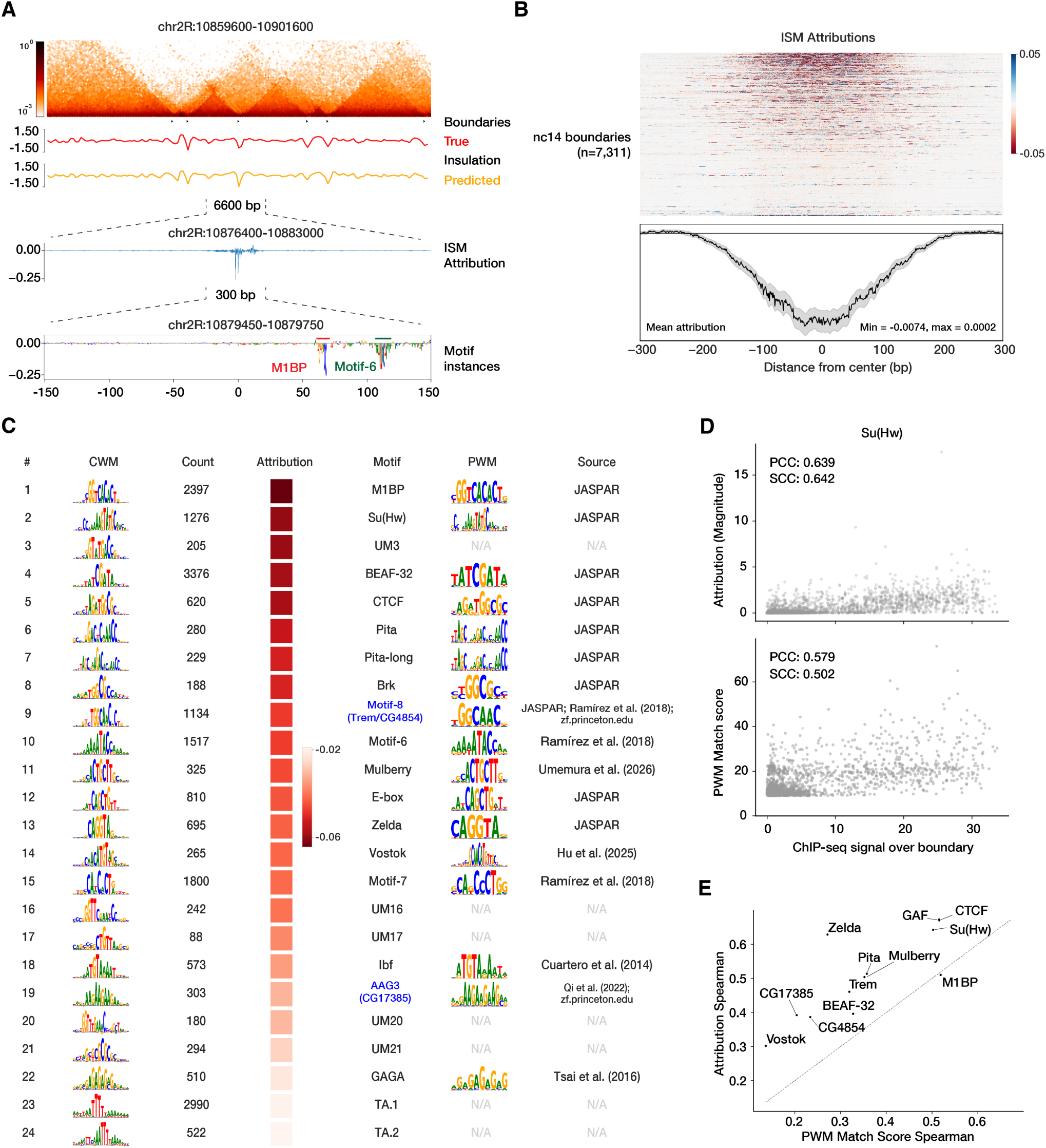
Sequence attribution analysis identifies the motifs underlying chromatin insulation. (**A**) Base-pair resolution Domino attributions calculated using *in silico* mutagenesis (ISM). For a representative 40.2 Kb held-out region, attributions are shown for the 6.6 Kb region centered on a boundary, with a central 300 bp zoomed to visualize individual nucleotide insulation attributions. Negative attributions indicate sequence elements whose presence decreases the predicted score, thereby contributing to stronger local insulation and driving boundary formation, while positive attributions designate sequence features that weaken insulation. (**B**) ISM attribution profiles for the 600 bp sequences centered on 7,311 boundaries identified in the nc14 embryo. Top: Each boundary is represented by two rows (forward and reverse complement). Bottom: Mean per-base-pair attribution across all boundaries; shaded area represents ±2 standard errors of the mean, smoothed with a Gaussian 1D filter (*σ* = 2). (**C**) Motifs with negative attributions discovered *de novo* by Domino, ranked by average attribution. Each row displays the contribution weight matrix (CWM) from TF-MoDISco^31^, seqlet count, average attribution score, and the best match to known motifs (motif name, positional weight matrix (PWM) logo, and source of the motif). (**D**) Scatterplots compare Su(Hw) ChIP-seq signal against attribution magnitude (top) and PWM match score (bottom). PCC: Pearson correlation coefficient; SCC: Spearman correlation coefficient. (**E**) Domino attributions have better correlation with binding signal at boundaries than motif match scores. Comparison across 12 architectural factors with available embryonic ChIP-seq data. The scatterplot shows Spearman correlation of ChIP-seq signal with attribution (*y*-axis) and motif PWM match score (*x*-axis). GAF was evaluated at boundaries with GAGA motif instances; Trem and CG4854 were evaluated at boundaries with Motif-8 (see **Fig. 4** below).

Because Domino is a predictive model that uses a wide sequence context, we reasoned that its attributions would better capture functional factor activity than standard motif scanning^46^. To test this, we benchmarked both metrics against embryonic ChIP-seq data for 12 architectural factors (**Table S1**), using binding intensity as a proxy for factor activity. Remarkably, Domino attributions better correlated with ChIP-seq-measured binding intensity for 11 of the 12 factors (**Fig. 2D,E**), confirming that the model captures quantitative variation in functional factor activity rather than merely identifying motif occurrences.

Crucially, Domino also discovered insulation motifs for which no DNA-binding proteins have been previously identified. While a subset of them matched motifs previously reported as associated with promoters or boundaries^14,35,47^, others represented entirely novel sequences with no clear literature references (**Fig. 2C**). We anticipate that these uncharacterized motifs correspond to previously undescribed insulator factors in *Drosophila*. Below, we focus specifically on two such motifs, Motif-8 and AAG3, which enabled us to characterize the novel insulation factors CG4854, Trem and CG17385. Thus, Domino systematically uncovered known and discovered new sequence-specific factors with insulatory activity.

### Domino predicts context-specific insulation activity of individual motif instances

While Domino enables the systematic characterization of insulation motifs through genome-wide modeling, an additional strength of interpreting this sequence-to-function framework lies in its ability to reveal the functional status of individual motif instances. Because the model leverages a wide sequence context, it can distinguish whether a specific motif instance is actively contributing to boundary formation or remaining silent. Indeed, we found that two identical copies of the core Motif-8 embedded within two separate canonical boundaries, *nhomie* and *homie*, received completely different functional attributions from the model. While the Motif-8 instance in *nhomie* was predicted as active, the identical sequence in *homie* had zero attribution (**Fig. 3A**). In contrast, neighboring Su(Hw) motifs received strong attributions in both contexts, consistent with the recent report that Su(Hw) drives insulation activity in both *nhomie* and *homie*^48^. This divergence indicates that identical core motif sequences can harbor distinct functional activities dictated by broader sequence context.

**Figure 3.**
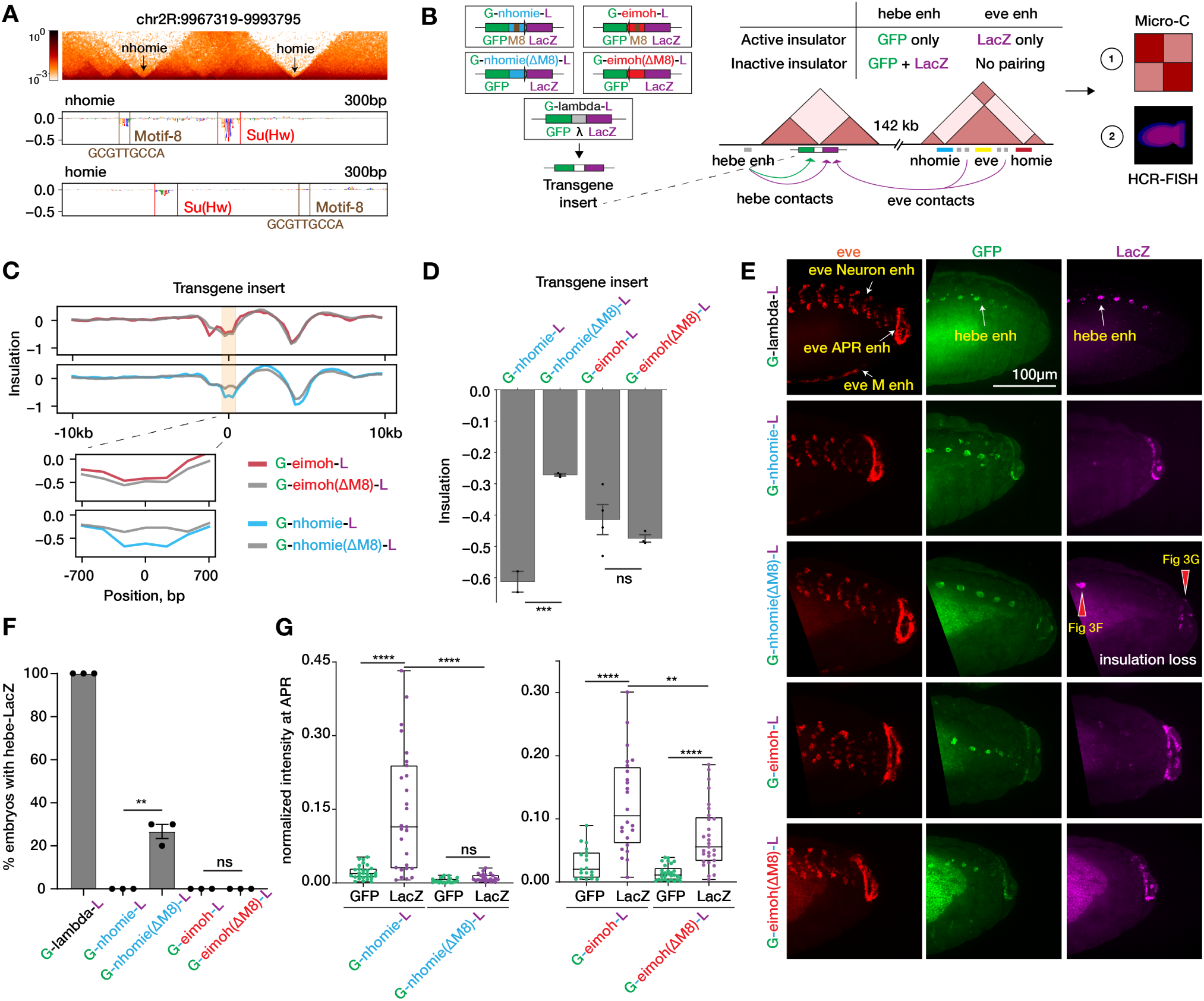
Domino predicts context-specific insulation activity of individual motif instances. (**A**) Context-specific attributions at the *nhomie* and *homie* boundaries. Domino identifies a Motif-8 instance at *nhomie* with strong negative attribution (predicted contribution to insulation) and an instance at *homie* with the identical core sequence (highlighted in brown) but no attribution. In contrast, the neighboring Su(Hw) motif (highlighted in red) has high insulation attribution in both boundaries. **(B–G)** Experimental validation of context-specific Motif-8 requirements from (A). **(B)** Transgene reporter system at the *hebe* locus. One of five dual-reporter (*GFP*, *LacZ*) transgene constructs (left) was inserted near *hebe*, 142 Kb upstream of the *eve* locus. Integrated constructs included wild-type and Motif-8 mutants (ΔM8) of *nhomie* and *homie* (inverted, “*eimoh*”) and *lambda* fragment control. Insulator activity was assessed using Micro-C (C–D) and HCR-FISH (E–G). HCR-FISH measured insulation at the transgene through blocking of the proximal *hebe* enhancer (E, F) and through long-range interactions between the transgene reporters and the endogenous *eve* APR (anal plate ring) enhancer (E, G). **(C)** Micro-C insulation tracks at the transgene insertion site. Mutation of Motif-8 results in a substantial reduction of insulation at *nhomie* but not *homie*. **(D)** Quantification of insulation shown in (C). ***, *p <* 0.001, one-sided *t*-test. **(E)** HCR-FISH for *eve*, *GFP* and *LacZ* in embryos. Active enhancer locations indicated. Red arrows highlight abnormal *LacZ* expression driven by the local *hebe* enhancer (indicating reduction of insulation) and reduced *LacZ* expression driven by the distal *eve* APR enhancer (indicating reduction of pairing, as a proxy of insulatory activity) in the *nhomie* Motif-8 mutant. M, mesoderm. **(F)** Percentage of embryos with *hebe* enhancer-driven *LacZ* expression. **, *p <* 0.01, non-parametric unpaired *t*-test. Each dot represents one replicate consisting of *n* = 10 embryos. The *nhomie* Motif-8 mutant (but not the *homie* mutant) has a significant breakdown of enhancer-blocking activity, allowing the local enhancer to bypass the boundary and activate the reporter. **(G)** Quantification of normalized *GFP* and *LacZ* expression in the embryonic region with active *eve* APR enhancer. The *nhomie* Motif-8 mutant (but not *homie* mutant) has reduced long-range reporter activation, indicating reduced pairing with the endogenous *homie* and *nhomie*. **, *p <* 0.01, ****, *p <* 0.0001, paired two-tailed *t*-test. Sample sizes: *nhomie*, *homie*, and *nhomie* ΔM8 mutant, each *n* = 30; *homie* ΔM8, *n* = 26.

To experimentally validate these context-specific predictions *in vivo*, we used a site-specific dual-reporter (*GFP*, *LacZ*) transgene integration system at the *hebe* locus, located 142 Kb upstream of the endogenous *nhomie–eve–homie* locus (**Fig. 3B, Fig. S3, Methods**). We integrated five unique boundary constructs, including wild-type and Motif-8 mutant (ΔM8) variations of *nhomie* and *homie* (inverted; “*eimoh*”), together with a neutral *lambda* fragment control, and evaluated structural and transcriptional readouts using high-resolution Micro-C and HCR-FISH. Consistent with Domino computational predictions, Micro-C profiling revealed that mutation of the Motif-8 site resulted in a substantial, localized reduction of insulation at the *nhomie* transgene insert, whereas mutating the same motif in the *homie* element produced no detectable change in activity (**Fig. 3C,D**). We next characterized the downstream transcriptional consequences of these perturbations using HCR-FISH. In the baseline control condition (*lambda*), the proximal *hebe* enhancer activated both flanking reporters, whereas wild-type *nhomie* and *homie* fragments robustly blocked *LacZ* activation, indicating insulatory activity (**Fig. 3E,F**). Strikingly, the Motif-8 mutation (ΔM8) caused significant enhancer bypass and an emergence of *hebe*-driven *LacZ* expression in up to 30% of *nhomie* mutant embryos, while *homie(*Δ*M8)* embryos remained fully insulation-competent (**Fig. 3E,F**). Furthermore, quantitative analysis of expression presumably driven by long-range regulatory interactions with the endogenous *eve* anal plate ring (APR) enhancer demonstrated that deletion of Motif-8 compromised distal reporter activation uniquely within the *nhomie* sequence context (**Fig. 3E,G**). Together, these findings demonstrate that Motif-8 is a strict, context-dependent requirement for physical insulation and enhancer-blocking at *nhomie* but not *homie*, confirming the unique capacity of Domino to decode functionality of individual motif instances across the genome.

### Domino guides the discovery of the zinc-finger proteins Trem, CG4854, and CG17385 as novel chromatin insulation factors

The systematic characterization of insulation motifs through our genome-wide predictive modeling provided a unique opportunity to identify previously uncharacterized sequence-specific insulation factors. Given that many known *Drosophila* architectural factors, including CTCF, Su(Hw), Pita, M1BP, Zelda and Mulberry, are Cys_2_His_2_ (C2H2) zinc finger (ZF) proteins^49^, we hypothesized that the uncharacterized insulation motifs identified by Domino may similarly be recognized by previously undescribed C2H2 ZF DNA-binding factors. To test this hypothesis, we leveraged an online resource that predicts DNA-binding motifs for C2H2 zinc-finger arrays using a machine learning framework trained on bacterial one-hybrid (B1H) data^50,51^. By screening our motifs against DNA-binding motif predictions for all four-zinc-finger arrays of C2H2 ZF domains in the set of *Drosophila* transcription factors, we successfully identified high-confidence protein candidate matches for two primary uncharacterized motifs.

We first investigated Motif-8, one of the strongest motifs conferring insulation in our analysis, which we had also identified as a context-specific determinant of insulation at the *nhomie–eve–homie* locus. Screening the Domino-derived Motif-8 against the predicted C2H2 ZF motifs nominated the uncharacterized protein CG4854 as the top significant match (**Fig. 4A, Supplementary File S2**). Because transcription factor paralogs frequently preserve DNA-binding specificities, we also evaluated *trade embargo (trem)*, a paralog of *CG4854* encoded in close proximity and likely sharing the same regulatory system (**Fig. S4A,B**). CG4854 and Trem both contain a zinc finger-associated domain (ZAD) linked to insulation (**Fig. S4A**)^49^. To determine if these factors indeed bind to Motif-8 *in vivo*, we analyzed public embryonic ChIP-seq datasets for both proteins^52^. *De novo* motif discovery independently recovered Motif-8 from both CG4854 and Trem binding peaks (**Fig. 4B,C, Fig. S4C, Methods**). Consistent with this finding, both factors had significantly higher ChIP-seq signal at genomic boundaries with strong negative Domino Motif-8 attributions than at other boundaries or control regions (**Fig. 4D**). More broadly, Trem and CG4854 ChIP-seq peaks, which significantly overlapped with one another across the genome, were frequently localized at boundaries (**Fig. 4C,E**). These data suggested that Motif-8 is bound by CG4854 and Trem to confer insulation. To validate this experimentally, we performed embryonic Micro-C profiling in *CG4854* and *trem* knockout (KO) strains (**Fig. 4F-H, Fig. S4D-F, Methods**). Loci bound by either factor exhibited a significant loss of insulation in KO embryos relative to wild-type controls (**Fig. 4G,H**). Both factors were preferentially expressed in the early embryo and adult ovaries, pointing to a specialized role in early development (**Fig. S4G**). Together, this analysis confirms CG4854 and Trem as sequence-specific insulation factors in the embryo.

**Figure 4.**
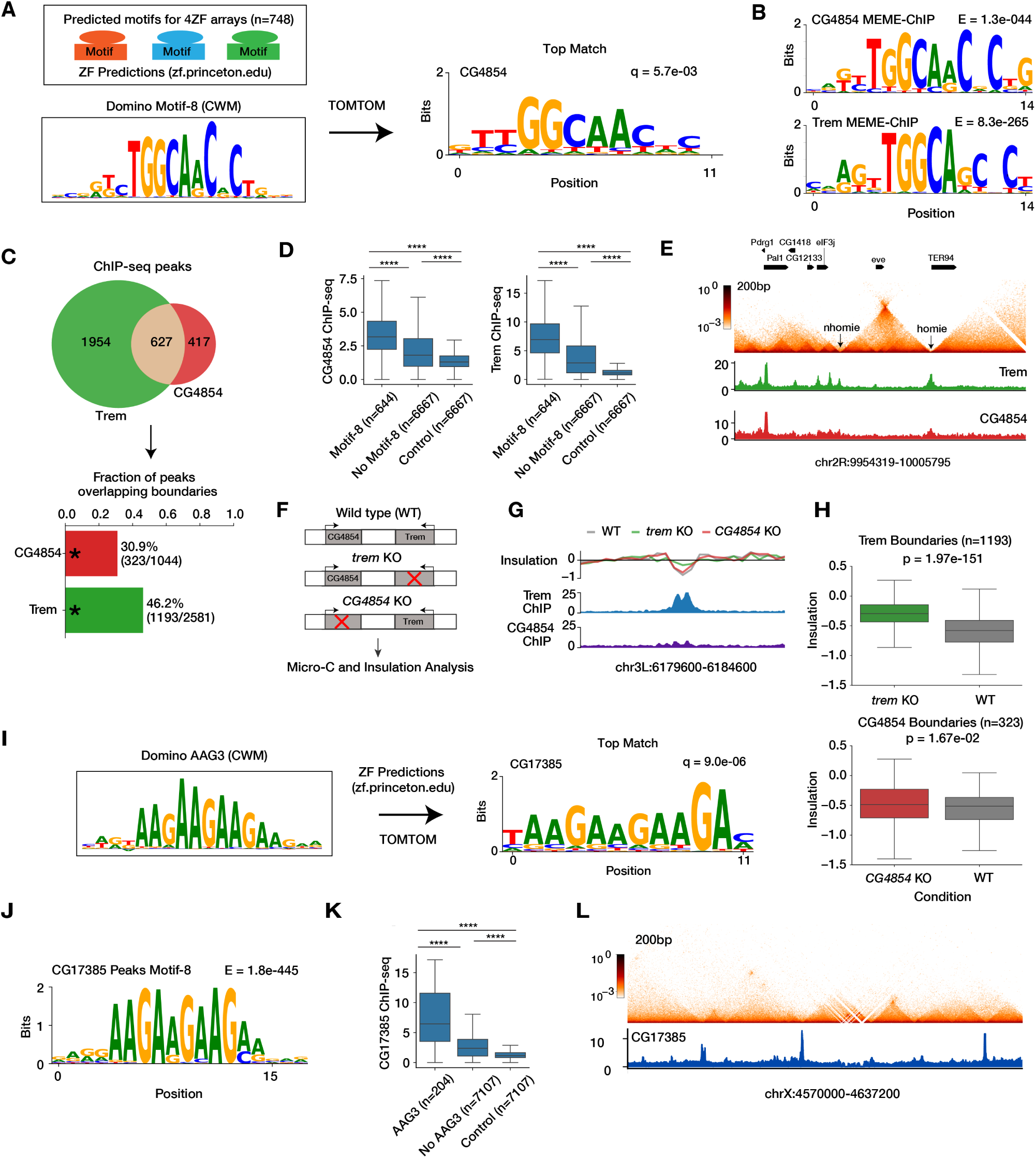
Domino identifies the zinc-finger proteins Trem, CG4854, and CG17385 as novel chromatin insulation factors. (**A**) Identification of CG4854 via motif matching. The Domino-derived Motif-8 motif (left) was compared against predicted motifs for 4-zinc-finger arrays^50,51^, identifying CG4854 as the top match (**Supplementary File S2**). (**B**) Motif discovery from public CG4854 and Trem ChIP-seq data^52^. CG4854 and Trem are paralogous zinc-finger proteins. The most significant motifs identified within Trem and CG4854 ChIP-seq peaks using MEME-ChIP^53^. The E-value represents the statistical significance of motif enrichment. (**C**) Trem and CG4854 binding co-localization. Venn diagram of Trem and CG4854 ChIP-seq peaks (top) and frequency of their overlap with the set of 7,311 identified embryonic boundaries (bottom). Significance was assessed with a permutation test; *, *p <* 0.0001. (**D**) Comparison of CG4854 (left) and Trem (right) ChIP-seq signal intensity between boundaries with Motif-8 attribution, boundaries without Motif-8 attribution, and non-boundary control regions. Significance was assessed using a one-sided Wilcoxon rank-sum test; ****, *p <* 0.0001. (**E**) Embryonic Micro-C and CG4854 and Trem ChIP-seq tracks for a *nhomie–eve–homie* genomic locus illustrating co-localization of insulation and CG4854 and Trem binding. (**F**) Knockout (KO) of *trem* or *CG4854* (encoded by adjacent genomic loci) was performed using a piggyBac insertion. Micro-C profiling was performed for the wild-type (WT), *trem* KO, and *CG4854* KO, followed by insulation analysis. **(G–H)** Micro-C analysis following *trem* and *CG4854* KO. (**G**) Example 5 Kb region with Trem binding showing insulation loss in *trem* KO vs. WT. (**H**) Genome-wide insulation comparison between *trem* KO and WT Trem-bound boundaries (*n* = 1193, top) and between *CG4854* KO and WT CG4854-bound boundaries (*n* = 323, bottom). Paired *t*-test. Boxplots: center line, median; box limits, upper and lower quartiles; whiskers, 1.5× interquartile range; outliers not shown. (**I**) Identification of CG17385 via motif matching. The Domino-derived AAG3 motif (left) was compared against predicted motifs for 4-zinc-finger arrays^50,51^, identifying CG17385 as the top match (**Supplementary File S2**). (**J**) Motif discovery from public CG17385 ChIP-seq data^52^. The most significant motif identified within CG17385 ChIP-seq peaks. The E-value represents the statistical significance of motif enrichment. (**K**) Comparison of CG17385 ChIP-seq signal intensity between boundaries with AAG3 attribution, boundaries without AAG3 attribution, and non-boundary control regions. Significance was assessed using a one-sided Wilcoxon rank-sum test; ****, *p <* 0.0001. (**L**) Embryonic Micro-C and CG17385 ChIP-seq tracks for a representative genomic locus illustrating co-localization of insulation and CG17385 binding.

We next examined the AAG3 motif^47^, which exhibited strong negative functional attribution across our embryonic boundaries but lacked a known binding factor. Matching the AAG3 motif against the C2H2 ZF motif predictions revealed that the top significant candidate was the uncharacterized protein CG17385 (**Fig. 4I, Supplementary File S2**). The expression profile of CG17385 indicated a preferential role in the early embryogenesis (**Fig. S4G**). *De novo* motif discovery from public embryonic CG17385 ChIP-seq data^52^ retrieved the AAG3 motif as the top enriched motif (**Fig. 4J, Methods**). Furthermore, genomic boundaries with strong negative Domino AAG3 attribution had a significant enrichment of CG17385 ChIP-seq signal compared to other boundaries or control regions (**Fig. 4K,L**), establishing CG17385 as a novel sequence-specific boundary-associated factor in the embryo.

Thus, Domino helped us identify three novel insulation factors from the C2H2 ZF family: CG4854, Trem and CG17385.

### A comprehensive motif grammar of chromatin insulation in the *Drosophila* embryo

Having demonstrated that Domino captures context-specific motif contributions to insulation genome-wide, we leveraged our high-resolution embryonic dataset to comprehensively characterize the global insulation landscape. Filtering for boundaries with at least one active motif (total attribution ≤ −0.25) across all 24 motifs (*n* = 4, 289 boundaries; **Methods**), K-means clustering identified 30 distinct clusters spanning established and new promoter-associated, developmental, and other architectural elements (**Fig. 5A, Fig. S5A,D-F**). Notably, while boundaries without an active motif had weaker insulation, Domino predicted their insulation profiles with an accuracy on par with motif-containing boundaries, indicating their insulation is still sequence-dependent (**Fig. S5B,C**); characterizing such boundaries and exploring the underlying sequence determinants could be an important target for future studies. The motif-based clusters correlated strongly with experimental *in vivo* binding data, showing robust factor-specific ChIP-seq enrichment across clustered boundaries (**Fig. 5A,B, Fig. S5G**). The landscape included large promoter-associated clusters (e.g. driven by M1BP and BEAF-32) with strong insulation near transcription start sites (TSSs), alongside a diverse array of clusters localized farther from TSSs (**Fig. S5F**). These distal boundaries are driven by developmental factors (Zelda, GAF), classic structural insulators (Su(Hw), CTCF, Pita, Ibf), new factors described by us and others (CG4854, Trem, Mulberry, Vostok, CG17385), and completely novel motifs that likely correspond to undiscovered binding proteins. Together, these distinct clusters establish a comprehensive classification of the insulation repertoire in the *Drosophila* embryo.

**Figure 5.**
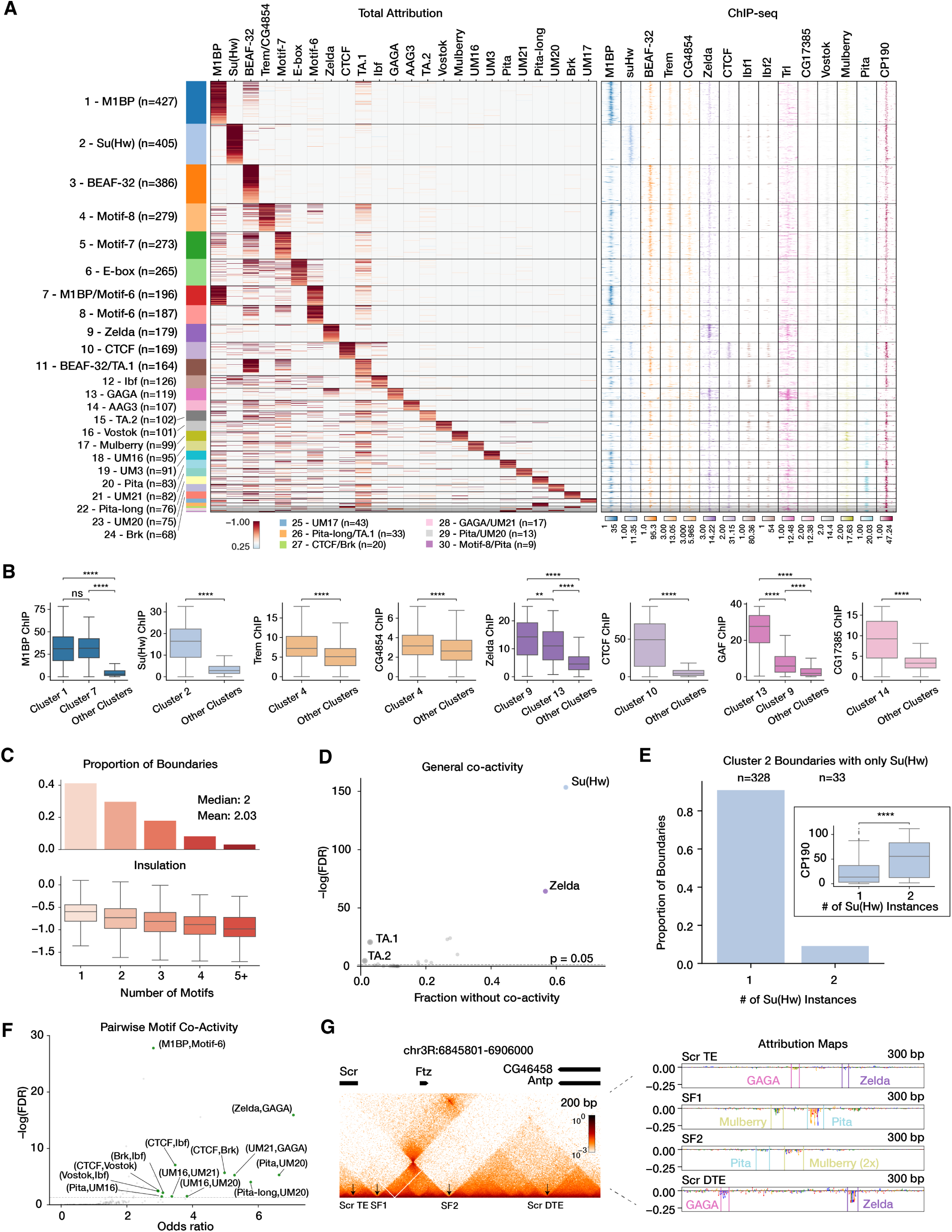
A comprehensive motif grammar of chromatin insulation in the *Drosophila* embryo. (**A**) Clustermap of the insulation motif repertoire. Attributions for all 24 motifs (columns) across boundaries containing at least one active site (*n* = 4, 289, rows). Motifs are defined as active if their total attribution is *<* −0.25. Clusters are sorted by size, with rows within each cluster ranked by total attribution. Tornado plots show ChIP-seq occupancy signal for corresponding factors across 3.2 Kb regions centered on the boundaries. **(B)** ChIP-seq signal enrichment at selected clusters. Comparison of insulator factor ChIP-seq signal intensity between boundaries in the cluster corresponding to the factor and boundaries from all other clusters. Along with new factors Trem, CG4854, and CG17385, five other representative insulator-associated factors are selected. Zelda binding is enriched in cluster 9 with Zelda motif and cluster 13 with GAGA motif and some presence of Zelda motif. GAF binding is enriched in cluster 13 with GAGA motif and in cluster 9 without GAGA but with Zelda motif. Significance is assessed using a two-sided Wilcoxon rank-sum test; ****, *p <* 0.0001. Boxplots: center line, median; box limits, upper and lower quartiles; whiskers, 1.5× interquartile range; outliers not shown. **(C)** Active motif density and insulation distribution. Distribution of the number of active motifs per boundary (top). Relationship between motif count and insulation (bottom). Boxplots showing distributions of local insulation scores across boundaries grouped by the number of active motifs per boundary. Boxplots: center line, median; box limits, upper and lower quartiles; whiskers, 1.5× interquartile range; outliers not shown. **(D)** Exclusive and promiscuous motifs. For each motif, *x*-axis value shows the fraction of boundaries with the motif which do not have other active motifs. Significance is assessed via a two-sided Fisher’s exact test, corrected for multiple hypothesis testing (false discovery rate, FDR). Gray dots indicate all motifs with frequent co-activity (*<* 0.03 fraction without co-activity). Other colored dots (colored by motif-associated cluster from (A)) indicate motifs with infrequent co-activity (*>* 0.35 fraction without co-activity). **(E)** Most Su(Hw)-associated cluster 2 boundaries contain a single copy of Su(Hw) motif. Barplot showing fraction of cluster 2 boundaries with the specified number of active Su(Hw) motif instances. Inset: Boxplots showing the distribution of CP190 ChIP-seq signal in cluster 2 depending on the number of active Su(Hw) motif instances. Significance is assessed using a two-sided Wilcoxon rank-sum test; ****, p < 0.0001. One boundary which contained more than two Su(Hw) instances was omitted. Boxplots: center line, median; box limits, upper and lower quartiles; whiskers, 1.5× interquartile range; outliers not shown. **(F)** Pairwise motif co-activity. The odds ratio for co-occurrence of two motifs in a boundary. Significance of enriched or depleted co-occurrence for motif pairs is assessed via a one-sided Fisher’s exact test, corrected for multiple hypothesis testing. Green dots indicate all significant motif pairs with OR > 2.5 that did not include TA-rich motifs. **(G)** Attribution analysis of boundaries and tethering elements (TE) at the *Scr* and *Ftz* loci. Left: The contact map at this region that contains previously described boundaries SF1 and SF2 and a TE near *Scr* promoter that loops with a distal TE (DTE)^16^. Right: Domino attribution maps for 300 bp regions at SF1, SF2, *Scr* TE, and *Scr* DTE. Attribution maps highlight co-activity of Pita and Mulberry motifs in SF1 and SF2 and co-activity of GAGA and Zelda in *Scr* TE and DTE.

Beyond global boundary classification, we systematically analyzed motif co-activity. Surprisingly, motif-containing boundaries had, on average, only two active motifs, and more than 40% of boundaries had a single active motif, although an increased density of active motifs scaled progressively with stronger local insulation (**Fig. 5B**). This low motif density indicates that individual boundary motif grammar is amenable to comprehensive characterization. Systematic co-activity analysis revealed distinct extremes: while AT-rich (TA) motifs frequently co-occurred with other motifs mostly at promoter-associated boundaries, the Su(Hw) and Zelda motifs exhibited particularly low overall co-activity, rarely appearing alongside other active motifs within the same boundary (**Fig. 5C**). Although these exclusive motifs generally conferred weaker insulation than other motifs, their insulation levels were significantly stronger than background (**Fig. S5E**). The Su(Hw) motif was the most exclusive, exhibiting a strong depletion of other architectural factors with the notable exception of CP190, consistent with its established recruitment mechanism (**Fig. 5A**)^37^. Furthermore, the vast majority of Su(Hw) boundaries had only a single active Su(Hw) motif instance; the rare boundaries with two active instances had increased CP190 binding, consistent with a dose-dependent co-activity (**Fig. 5E**). Intriguingly, Su(Hw) also displayed the most significant tendency to occupy consecutive genomic boundaries, suggesting a specialized role in local chromatin looping across a distinct set of boundaries (**Fig. S5H**)^48^.

We next focused on systematic analysis of pairwise motif co-activity (**Fig. 5F, Fig. S5I**). For example, M1BP and Motif-6 co-occurred across genomic boundaries significantly more frequently than expected by chance, consistent with their previously described association with promoters^14^. GAGA and Zelda motifs also showed significant co-occurrence (**Fig. 5F, Fig. S5I**); furthermore, Zelda binding was enriched in the GAGA-associated boundary cluster despite having Zelda motif only in a subset of those boundaries, and GAF (GAGA-associated factor encoded by *Trithorax-like* (*Trl*) gene) binding was enriched in the Zelda-associated boundary cluster despite lacking GAGA motif (**Fig. 5A,B**), suggesting joint Zelda and GAF recruitment to these boundaries. As a functional illustration, we detected both active motifs within the previously characterized *Scr* tethering elements TE and DTE in the *ftz* locus, which interact through GAF-driven tether-tether pairing^16^ (**Fig. 5G**) Although both *Scr* TE and DTE exhibited relatively weak insulation and fell below the threshold for our main boundary list, they nonetheless conferred noticeable insulation. By performing targeted sequence attribution analysis at these specific loci, we resolved active instances of both GAGA and Zelda motifs (**Fig. 5G**), further illustrating the interaction between Zelda and GAF.

In sum, our systematic dissection of motif attributions revealed a comprehensive motif grammar of chromatin insulation in the *Drosophila* embryo.

### Distance– and orientation-dependent insulation motif synergy

Given that many insulation motifs exhibited statistically enriched patterns of co-occurrence, we next investigated whether specific motifs may be synergistic in driving insulation. We quantified synergy *in silico* as the relative change in predicted insulation when both motifs are simultaneously present compared to the sum of their individual sequence predictions (**Fig. 6A, Methods**). For this, we used Domino predictions for synthetic sequences obtained by embedding single or multiple motifs into neutral endogenous genomic background sequences lacking insulation. We systematically computed these synergy scores for all pairwise combinations of the 20 motifs with most frequent negative attribution across all possible relative orientations, orders, and distances ranging from 1 to 150 bp.

**Figure 6.**
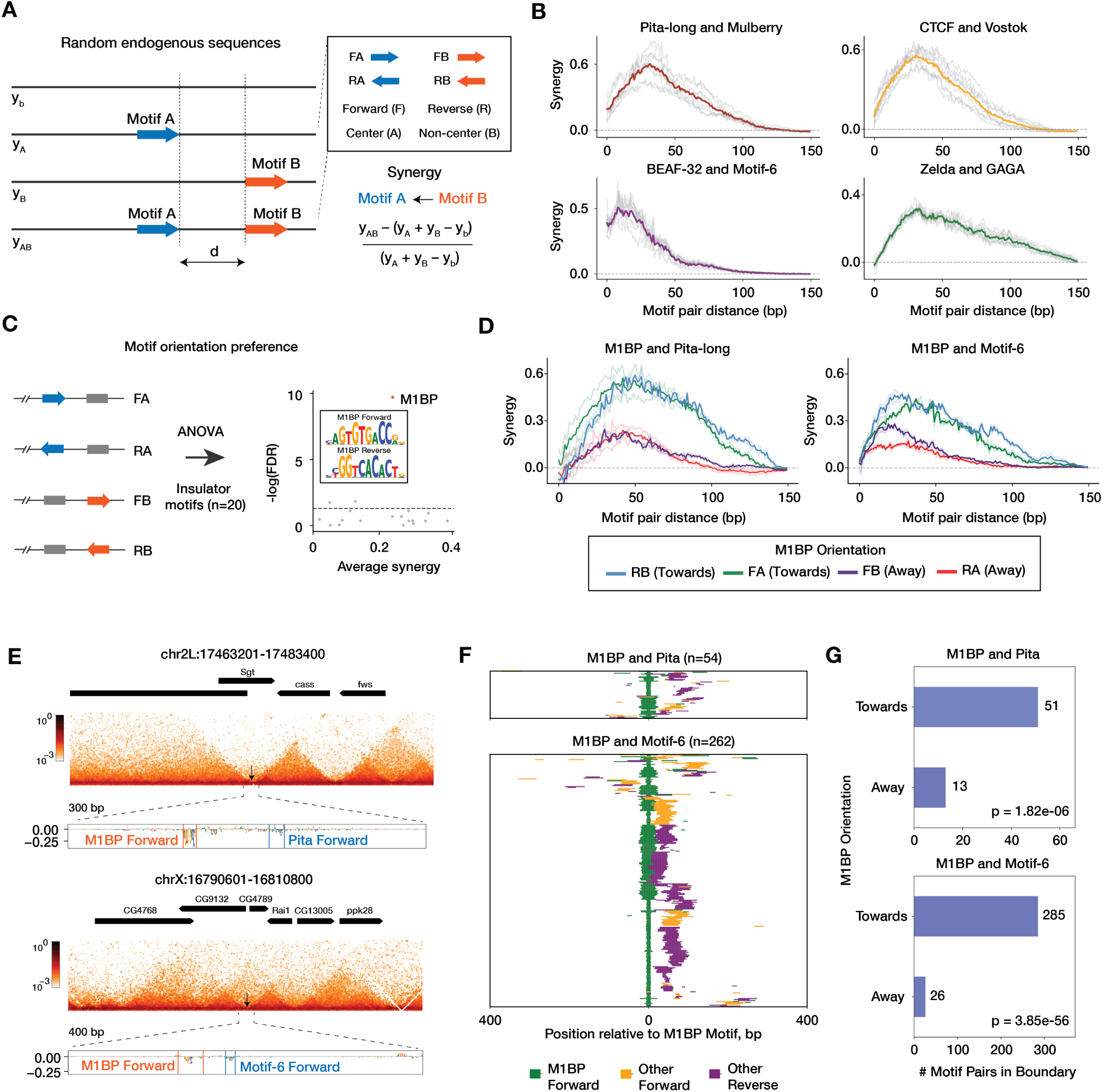
Distance– and orientation-dependent synergy between insulation motifs. (**A**) Calculation of motif synergy. Synergy is defined as the relative difference between the predicted insulation of a sequence containing both motifs and the sum of activity of each motif individually. Motifs are synthetically embedded into sequences sampled from endogenous genomic regions lacking insulation. **(B)** Distance-dependent synergy. Synergy tracks for selected motif pairs across distances 1–150 bp. Gray lines show individual tracks for all possible relative orientations and positions; the colored line represents the mean synergy across all configurations. **(C)** Systematic evaluation of orientation and order preferences across the 20 most frequent insulation motifs reveals M1BP as the most significant orientation-dependent motif. For each of the four relative orientations of a motif *X* (colored arrow), a distribution of synergy scores with every other motif *Y* (gray bar; tested for both orientations) was computed as the average of the top 20 synergy scores between *X* and *Y* for *d* between 1 to 150 bp. For each group, ANOVA was performed to test if the motif *X* exhibited an orientation preference. **(D)** Synergy tracks for M1BP and Pita-long motifs (left) and M1BP and Motif-6 motifs (right) across 1–150 bp. Blue and green tracks represent configurations where M1BP is oriented toward the partner motif; purple and red tracks represent configurations where M1BP is oriented away. Thicker curves show the average between the two respective curves of the specified configuration (as illustrated in (C)), where the partner of M1BP can be in the forward or reverse orientation. **(E)** Representative loci. Contact maps and corresponding Domino attribution maps for boundaries containing M1BP paired with Pita (top) or Motif-6 (bottom). Attribution maps highlight M1BP instances oriented toward their respective partners. **(F)** Genome-wide distribution of motif co-occurrence. Heatmaps show the relative positions and orientations of M1BP and its partners (Pita, combining both Pita and Pita-long motifs, top; Motif-6, bottom) across all boundaries with identified co-activity. All examples are anchored on the M1BP instance in the forward orientation. **(G)** Significant association between M1BP orientation and its relative order to Pita (top) and Motif-6 (bottom) instances withing boundaries genome-wide (one-sided Fisher’s exact test). Barplots show the number of genomic co-occurrences in each orientation configuration.

We first evaluated whether functional synergy exists between motif pairs and if it aligns with the endogenous grammar of real boundaries. *In silico* predictions revealed robust synergy across various motif combinations (**Fig. 6A,B, Fig. S6**), mirroring and extending the statistical patterns of co-activity discovered in our native boundary analysis (**Fig. 5F, Fig. S5I**). Crucially, the use of synthetic sequence backgrounds enabled us to systematically explore distance and orientation preferences independent of genomic confounding factors. By averaging synergy tracks across all eight relative configurations for each motif pair, we found that pairs characterized by frequent genomic co-occurrence exhibit distinct distance-dependent profiles (**Fig. 6B**). Certain motif pairs, for example BEAF-32 and Motif-6, displayed maximal synergy at ultrashort intervals (*<* 20 bp), establishing a hard syntax typical of direct protein-protein interactions^24,54^. Conversely, other pairs exhibited a more flexible, soft syntax spanning wider spatial intervals. For example, Pita-long/Mulberry (see **Fig. 5G** for an example) and CTCF/Vostok synergies showed a peak at approximately 40 bp, and Zelda and GAGA exhibited cooperative insulation across a broad window spanning 30 to 70 bp, corroborating our earlier observations of their cooperation. Rather than a direct physical contact, such soft syntax patterns suggest an indirect cooperativity mediated through local co-factor recruitment or the formation of higher-level multi-protein hubs.

Because motif synergy can also be dictated by spatial directionality, we systematically evaluated orientation and order preferences across the top 20 motifs (**Fig. 6C**). We found that M1BP was the most definitive orientation-dependent architectural factor. A detailed inspection of M1BP partner synergies revealed that this structural bias was driven primarily by its interactions with Pita and Motif-6. Specifically, insulation synergy was maximized only when the M1BP motif was physically oriented toward its neighboring partner motif (**Fig. 6D**). To determine whether these *in silico* orientation preferences reflect constraints of native boundary organization, we mapped the distribution of these multi-motif configurations across endogenous embryonic boundaries genome-wide. We observed a significant orientation-specific pattern: when M1BP co-occurs with Motif-6 or Pita, it is preferentially oriented toward its partner (**Fig. 6E–G**). Given that the M1BP protein features a ZAD domain, capable of mediating protein-protein interactions and implicated in insulation^49^, followed by a DNA-binding zinc-finger array, this strict orientation constraint presumably serves to position the M1BP interaction surface optimally relative to binding partners to facilitate directional chromatin insulation.

Together, these findings reveal spatial boundary syntax in the *Drosophila* genome, where motif synergy is controlled by distance, orientation, and order.

### Domino resolves a dynamic and complex chromatin insulation landscape in the *Drosophila* brain

Chromatin architecture is highly dynamic across developmental stages and tissues. To investigate how these dynamics are encoded at the sequence level, we applied our Domino framework to characterize tissue-specific insulation in the *Drosophila* larval and adult brain using public Micro-C datasets^19^ (**Fig. 7A**). This analysis revealed a substantially more extensive insulation landscape in neural tissues compared to the early embryo. While the majority of nc14 boundaries remained active in the larval and adult brains, a substantial number of additional genomic regions emerged as specialized insulators in the brain (**Fig. 7B,C**). Domino models trained independently on these neural Micro-C datasets achieved high predictive performance on held-out genomic regions (**Fig. S7**). This was particularly notable given that the brain datasets had lower sequencing coverage—271 million and 411 million contacts for larval and adult brain, respectively—than our integrated, ultra-large embryonic atlas with 1.9 billion contacts. Despite this difference in data depth, the models remained highly robust, allowing us to leverage model interpretation to systematically decode the sequence basis of tissue-specific insulation.

**Figure 7.**
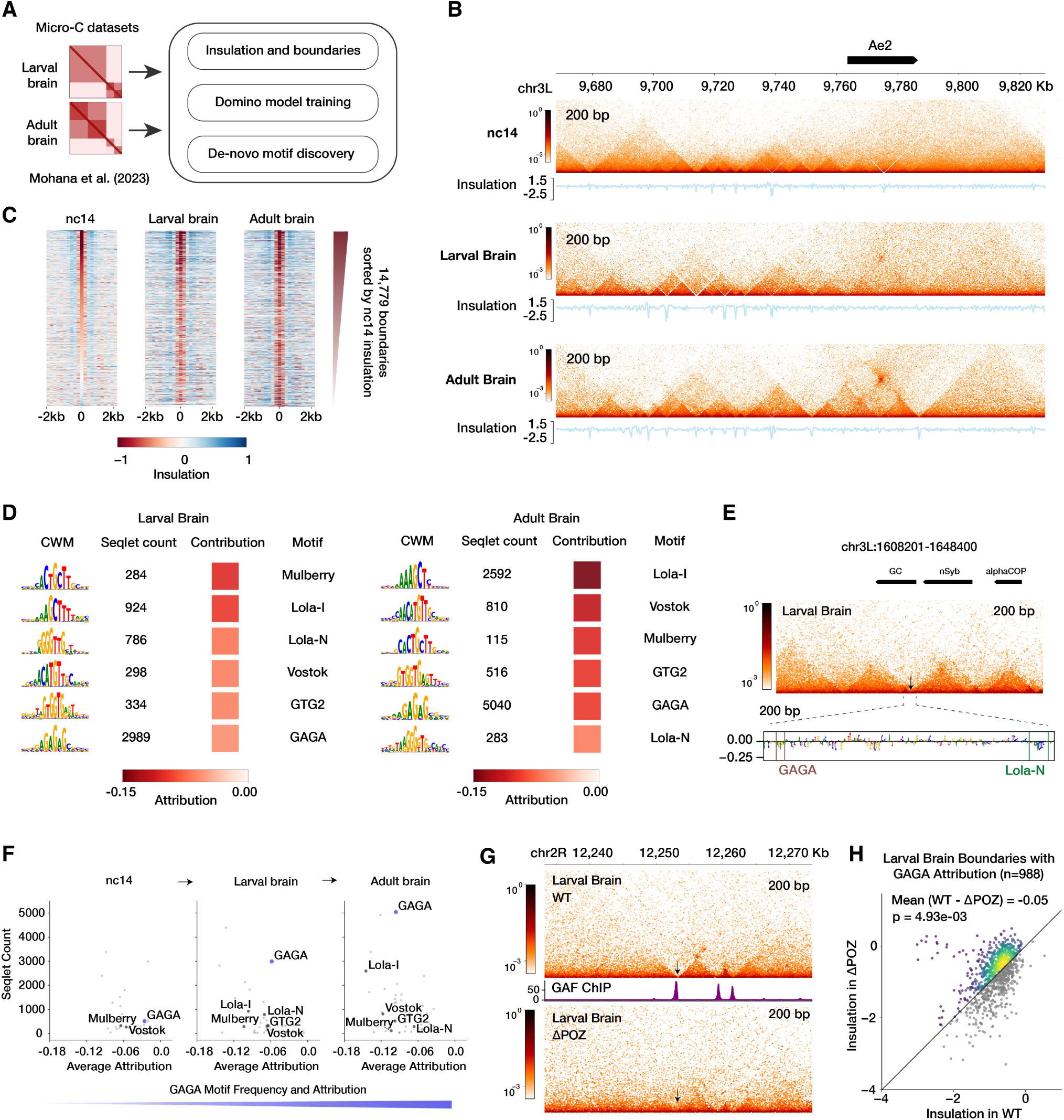
Chromatin insulation grammar in the *Drosophila* larval and adult brain. (**A**) Overview of Domino applied to Micro-C datasets from larval and adult brain^19^. **(B)** Temporal insulation dynamics at a representative locus flanking *Anion exchanger 2* (*Ae2*), a gene essential for neurogenesis and brain development. Micro-C contact maps and corresponding insulation tracks for an 160.2 Kb region showing a progressive increase of insulation from the nc14 embryo to the larval and adult brain. Genes other than *Ae2* that are present in this region are not shown. **(C)** Tornado plots show insulation scores at the union of all boundaries across nc14 embryos, larval and adult brain. Regions (4.2 Kb) are centered on boundaries and sorted by insulation strength in nc14. **(D)** Brain-specific insulation motifs. Selected motifs discovered *de novo* from boundaries that emerge or strengthen in the brain relative to the embryo. (Full list shown in **Fig. S8, S9, S10**.) Each row shows the CWM from TF-MoDISco, seqlet count, average attribution, and the best match to known motifs. **(E)** A representative boundary in the larval brain near a gene *neuronal Synaptobrevin* (*nSyb*). A contact map and the corresponding Domino attribution map for a boundary containing Lola-N and GAGA motifs. **(F)** Scatterplots showing average attribution and seqlet count for motifs across conditions. Motifs highlighted in (D) are labeled if they were identified as active. **(G)** Representative contact maps using published Micro-C data^19^ showing the loss of insulation at a GAGA-associated boundary in the ΔPOZ GAF mutant compared to wild-type (WT), along with GAF ChIP-seq in the larval brain^42^. **(H)** Genome-wide reduction of insulation in the ΔPOZ GAF mutant. Scatterplot shows insulation in WT vs. ΔPOZ GAF mutant across boundaries with nonzero GAGA motif attribution (*n* = 988). One-sided paired *t*-test. Colored points indicate boundaries with insulation loss in the mutant.

Our analysis identified a diverse set of insulation sequence features that increased in frequency and intensity compared to the nc14 embryo, bridging classic architectural factors with new ones (**Fig. 7E**, **Fig. S8, S9, S10**). Alongside established insulator elements, we discovered multiple motifs matching known factors previously not explicitly implicated in insulation, as well as novel uncharacterized motifs. For example, consistent with reports of specific GAF activity in the brain^29^, Domino revealed a marked increase in the frequency and attribution strength of the GAGA motif during the embryo-to-brain transition (**Fig. 7E,F**). To confirm a causal role for this factor, we analyzed public larval brain Micro-C data from GAF ΔPOZ mutants^19^. GAF mutation resulted in a significant genome-wide loss of insulation specifically at sites with active GAGA attributions, validating its requirement for boundary formation or maintenance (**Fig. 7G,H**). We also identified a prominent association of the Vostok motif with insulation, consistent with the recently described regulatory and architectural role of Vostok in the brain^42^. Intriguingly, our model interpretation successfully distinguished between the motifs of different isoforms of the behavioral regulator Lola. Domino identified the motif for Lola-I, an isoform recently shown to act as a pioneer and architectural factor driving RNA Polymerase II pausing and distal metalooping in the brain^55,56^. Remarkably, the model also identified distinct motifs corresponding to entirely different Lola isoforms, most prominently Lola-N. We found that the Lola-N motif, distinct from the Lola-I motif, significantly co-occurred with active GAGA sites specifically within neural boundaries (**Fig. 7E, Fig. S10**), nominating Lola-N as a new candidate architectural factor.

In sum, these findings demonstrate that Domino can successfully map tissue-specific insulation codes, revealing that the *Drosophila* brain uses a highly complex and specialized motif grammar to shape its unique 3D genome.

## Discussion

While the role of architectural proteins in global chromosome organization continues to be actively studied, the precise cis-regulatory code that dictates local chromatin insulation has remained elusive. Traditional motif-scanning methods merely register sequence presence without functional context, while recent deep learning methods optimized for full 2D contact-map prediction^22,23,57^ are often too complex or lack resolution to directly isolate local boundary rules. In this work, we developed Domino, a sequence-to-function deep learning framework that advances boundary analysis by combining ultra-deep *Drosophila* Micro-C data at 200 bp resolution with a targeted modeling strategy focused exclusively on the local insulation score rather than the entire contact matrix. Because the calculation of an insulation score inherently aggregates data using a weighted average over a sliding window, it effectively filters out experimental noise, improving the model’s ability to capture the underlying sequence signal. Furthermore, this design relies on the key insight that local insulation serves as a robust proxy for a wide array of interconnected chromosomal features, ranging from promoter and enhancer activity to local and distal looping. By leveraging this proxy, Domino evaluates the functional status of individual motif instances based on their local genomic environments, providing an interpretable and generalizable platform to dissect the rules governing three-dimensional genome architecture.

Our systematic analysis of embryonic boundaries revealed a remarkably concise grammar of structural organization, where motif-dependent genomic insulation comprises 59% of all insulation and is explained by a short list of only 24 motifs, with an average of just two active instances per boundary. Through this comprehensive profiling, Domino successfully recovered the motifs of most known *Drosophila* architectural factors, including the core CP190-recruiting proteins CTCF, Su(Hw), Pita, M1BP, Ibf, and BEAF-32^13,36–38,58–60^. A small fraction of the strongest boundaries are characterized by the combinatorial mixing and matching of multiple distinct motifs; this structural redundancy explains why global chromosome architecture is often robust to the experimental depletion of individual factors. Conversely, a substantial fraction of boundaries are driven by a single active motif, most notably Su(Hw) and Zelda. These exclusive sites typically track with weaker insulation levels, suggesting they define specialized local sub-TAD boundaries or localized tethering elements rather than major global domain walls. Beyond mapping these established elements, a paramount strength of the Domino framework lies in its predictive capacity to discover entirely new, uncharacterized insulation motifs. These novel motifs provide a valuable resource for the community, and future research will hopefully identify which specific transcription factors bind to these newly uncovered sites. Indeed, for two motifs identified by our model, we were able to use C2H2 zinc-finger DNA-binding specificity predictions to nominate their cognitive binding partners, the zinc-finger proteins CG17385, Trem, and CG4854, newly characterized factors conferring insulation in the early embryo.

Because Domino is a predictive sequence-to-function model, it can compute quantitative insulation scores for any arbitrary, user-defined DNA sequences. Assuming the framework has accurately captured the underlying endogenous sequence patterns during training, this predictive capability enables the systematic *in silico* dissection of spatial syntax using synthetic or semi-synthetic sequences. By computationally embedding motifs at various spatial configurations, we bypassed native genomic confounding factors to systematically uncover the rules governing distance– and orientation-dependent motif synergy. Most remarkably, this approach revealed that the prominent CP190-recruiting factor M1BP cooperates with neighboring motifs in a strict orientation-dependent manner. This distinct directional requirement is likely explained by specific biochemical mechanisms of conferring insulation, to be revealed in future studies.

Furthermore, the Domino framework can be readily deployed across different datasets, developmental stages, and tissues. By re-running Domino on public *Drosophila* Micro-C datasets from the larval and adult brain, we uncovered a comprehensive and highly dynamic neural motif grammar, demonstrating the model’s robust performance even when trained on data with substantially lower sequencing coverage than our embryonic map. This cross-tissue analysis identified a shifting regulatory landscape, including both conserved elements and new, brain-specific chromatin insulation motifs. This highlights the broader utility of sequence-to-function modeling in discovering novel tissue-specific architectural components.

Looking forward, several critical avenues remain to expand and enhance this paradigm. A major next step will be to incorporate multi-task learning frameworks that integrate other functional genomic modalities, such as transcription factor binding profiles and quantitative gene expression data, to model the direct interplay between 3D structure and transcriptional regulation. Additionally, future iterations of Domino can be extended beyond local insulation scores to explicitly capture higher-order features of chromosomal topology, including local enhancer-promoter and boundary looping, distal metaloops, and macro-scale trans-chromosomal interactions. Finally, applying this approach across organisms, including to more complex mammalian genomes, will test the universality and evolutionary conservation of the sequence rules of insulation and potentially provide a predictive toolkit for engineering synthetic human insulators and designing therapeutically customized 3D genomic topologies.

## Acknowledgments

We thank all members of the Pritykin lab for helpful discussions. This study was supported in part by NIH/NIAID grant DP2AI171161, Ludwig Institute for Cancer Research and Weill Cancer Hub East.

## Code Availability

The Domino software package is available at https://github.com/pritykinlab/domino.

## Methods

### Plasmid construction and transgenic fly lines

The WT Oregon-R strain is maintained and used in the Schedl laboratory at Princeton University. The *CG4854* and *trem* KO lines were obtained from Bloomington *Drosophila* Stock Center. The *CG4854* KO line (w^1118^; PBacWHCG4854^f02794^, Bloomington stock number 18586) contains a Piggibac insertion at *CG4854* gene body near promoter (3R:20,028,089..20,028,090). The *trem* KO line (w^1118^; PBacWHtrem^f05981^, Bloomington stock number 85711) contains a Piggibac insertion at *trem* gene body (3R:20,031,266..20,031,267). The expression of *trem* is completely depleted as described in previous study^61^. All lines were validated by genomic-prep and sequencing.

The dual reporters construct was described previously^62^. Briefly, each reporter contains the *eve* basal promoter (−275 to +106 bp relative to *eve* start site), either the LacZ (*eve-LacZ*) or EGFP (*eve-gfp*) coding region, and the *eve* 3’ UTR (+1300 to +1525 bp). These two reporters are divergently transcribed. Test fragments were then inserted between the two reporters. The fragments used here are: a 500 bp fragment from lambda phage DNA, a 367 bp wild-type homie fragment (CDEF)^63^, a 660 bp wild-type nhomie fragment, and homie and nhomie fragments with their Motif-8 sequences mutated. Exact sequences of the constructs are provided in **Supplementary File S1**.

The −142 kb attP landing site was described previously^64^. It contains two attP target sites for phiC31 recombinase-mediated cassette exchange (RMCE)^65^ and mini-white as a marker. RMCE can result in the insertion of the transgene in either orientation, and all four possible insertions of the transgenes can be recovered. All the transgenic flies were genomic-prepped and sequence validated.

### Micro-C library preparation

Micro-C data for nc14 embryos was generated as previously described^18^. Specifically, flies were placed into population cages for egg laying. Next, embryos were collected on yeasted apple juice plates in population cages for 4 hours, incubated for 12hr at 25°C, then subjected to fixation as follows. For st14 embryos, the procedure for collection was identical to nc14 but was performed overnight. Embryos were dechorionated for 2min in 3% sodium hypochlorite, rinsed with deionized water, and transferred to glass vials containing 5 mL PBST (0.1% Triton-X100 in PBS), 7.5 mL n-heptane, and 1.5mL fresh 16% formaldehyde. Crosslinking was carried out at room temperature for exactly 15min on an orbital shaker at 250rpm, followed by addition of 3.7 mL 2M Tris-HCl pH7.5 and shaking for 5min to quench the reaction. Embryos were washed twice with 15 mL PBST and subjected to secondary crosslinking. Secondary crosslinking was done in 10mL of freshly prepared 3mM final DSG and ESG in PBST for 45 min at room temperature with passive mixing. The reaction was quenched by addition of 3.7mL 2M Tris-HCl pH7.5 for 5min, washed twice with PBST, snap-frozen, and stored at −80°C until library construction. For 4 of the nc14 samples, crosslinking was performed using formaldehyde only and all the other samples used dual crosslinking (EGS/DGS).

Micro-C libraries were prepared as previously described^66^ with the following modifications. Embryos were incubated in 500 *µ*L TE–NP40 (2% NP40 in TE buffer containing 4 *µ*U/*µ*L micrococcal nuclease) at 4°C for 1 h, followed by the addition of 3.5 *µ*L of 1 M MgCl_2_ and 0.5 *µ*L of 1 M CaCl_2_ to activate micrococcal nuclease. Embryos were then be incubated at 16 °C for 45 min to fragment chromatin. Fragmented chromatin was end-labeled with biotinylated nucleotides by T4 PNK and Klenow fragments and ligated by T4 DNA ligase. Cells were subsequently lysed using proteinase K and 1% SDS, followed by phenol–chloroform extraction and ethanol precipitation. Purified DNA was sonicated to an average fragment size of 350 bp using a Covaris ME220. Finally, biotinylated libraries were purified with Streptavidin beads and barcoded, pooled, and subjected to paired-end sequencing on an Illumina NovaSeq platform (50 bp paired-end reads with an 8 bp index read).

### Micro-C data preprocessing

We used 19 high-resolution Micro-C sample datasets profiled from *Drosophila* embryos at the nc14 stage, including publicly available and newly generated data (**Table S1**). First, we used BWA-MEM^67^ to align the paired-end reads to the *Drosophila melanogaster* Release 6 reference genome (dm6)^68^ using the parameters –S –P –5 –M. Then, we used pairtools^69^ to find ligation junctions to form pairs, sort pairs, remove PCR duplicates, and select all pairs using the conditions where pair_type was at least one read was unique and the other was rescued (“UU” or “UR” or “RU”). Next, we generate a pairs file using the split command. Finally, we used pairix^70^ to index the resulting files and Cooler^71^ to generate contact matrices at 100 bp resolution. In total, 11 these Cooler file datasets were used for training our model; the remaining 8 Cooler files were left out to assess correlation between independent measurements from the same condition (referred to as biological replicates). For each set of Cooler files, the contact matrices were merged using the merge tool and balanced using the zoomify tool from the Cooler package to generate multi-resolution cooler (.mcool) maps.

For Micro-C in the *trem* and *CG4854* knock-outs (KO) and the corresponding wild-type (WT) control in stage 14 *Drosophila* embryos, we selected for uniquely mapping reads by filtering for MAPQ value ≥ 60 and also downsampled the Cooler maps for *CG4854* and WT to match the total read count of *trem* using the random-sample tool from Cooler^71^. Processing of these samples was otherwise the same as above.

To process the Micro-C samples for the different dual-reporter transgene constructs (see **Fig. 3** for variants), we modified reference genomes for data alignment for each strain. First, we identified the exact *hebe* insertion site using the sequence TCGGATTGGATCGGATGGGATCGGAAACGTTTGGAT using the dm6 reference genome (exact coordinates at chr2R:9,836,564-9,836,599); then transgene sequence constructs which included only the concatenated sequences of *GFP*, one of the five boundary elements, and *LacZ* were inserted directly downstream of this site (starting at chr2R position 9,836,600). Genomes were indexed using faidx command from SAMtools followed by index comand from BWA package^67^. We then aligned each mutant to its respective genome using the same pipeline performed for nc14 Micro-C.

For analysis of the brain data, we gathered 6 published^19^ high-resolution Micro-C datasets in wild-type larval and adult brain as well as larval brain where the POZ-domain of GAF was mutated (2 replicates each). Data were processed and merged into.mcool format by the authors of the original study and provided to us.

### Computation of insulation score

We used the FAN-C^72^ package to compute local insulation scores genome-wide on balanced Micro-C data at 200 bp resolution. Specifically, a local insulation score was computed for each 200 bp bin by using a sliding window of size 1 Kb along the contact matrix diagonal to aggregate the relative frequency of chromatin interactions crossing over a given locus.

After computing local insulation scores genome-wide, we used the boundary identification tool of FAN-C to call local minima of the insulation scores using a delta parameter of 2. Each local minima identified has an associated boundary score, with higher values denoting stronger insulation. Finally, to allow for direct comparisons of the relative insulation from region to region, we mean-scale normalized every raw insulation score for a 200 bp bin relative to the mean insulation across 4.2 Kb centered on the bin. This normalized score is termed “insulation score” or simply “insulation” in the paper (unless otherwise specified).

### Insulation data processing and filtering

We used FAN-C^72^ to call local minima from the insulation score track at 200 bp resolution as described previously. In total, we obtained 97,807 such examples, all 200 bp in length. To ensure the dataset represented sites genome-wide, we sampled an equal number of sites from examples that were not local minima. From this combined set, we then performed several filtering steps for quality control. First, all examples where the 4.2 kb insulation score neighborhood around each example contained at least one null, positive or negative infinity value were discarded. Examples where the aforementioned neighborhood were not within the bounds of a given chromosome were also removed. Finally, we filtered out all examples that overlapped with the modENCODE v2 blacklisted sites^73^ in *Drosophila* and sites that were present in the mitochondrial chromosome (chrM). Finally, we used the Pysam^74^ package to obtain the 6600 bp-long DNA sequences centered around each 200 bp example.

In total, our entire dataset consisted of (sequence, insulation) pairs. For each pair, an example consisting of the reverse complement of each original sequence with the same insulation value was also included. This resulted in a final dataset with 330,278 (sequence, insulation) pairs. Similar to the approach of the previously published model DeepSTARR^54^, we used sequences from the first and second half of chromosome 2R for testing (8.7%) and validation (8.7%) respectively; the remaining examples were used for training.

### Domino architecture

Domino (**D**NA-t**o**-Insulation-Score **M**odel for Analyzing **In**sulation of Chr**o**matin) is a convolutional neural network (CNN) that uses a one-hot-encoding (A = [1, 0, 0, 0], C=[0, 1, 0, 0], G=[0, 0, 1, 0], T=[0, 0, 0, 1]) of 6600 bp-long DNA sequence input to predict insulation at the center 200 bp bin. The architecture of Domino is similar to those used in sequence-to-function convolutional neural network models^24,54,75^. The model utilizes a series of convolutional blocks and dense layers that together form the root, trunk and head modules. Unless otherwise specified, each convolutional block consists of a one-dimensional (1D) convolution, followed by GELU activation and batch normalization.

The root module uses two convolutional blocks with 1D convolutions containing 512 filters of size 5 and 256 filters of size 5 respectively to identify sequence motifs using filters which scan along the input sequence. The stem consists of five convolutional blocks, each with 1D convolutions with 256 filters of size 5 followed by an average pooling layer of size 2. We refer to such blocks as the “intermediate stem blocks.” Residual connections were added between each of these blocks. Following the intermediate stem blocks, we added an additional residual block containing a 1D convolution with 256 filters of size 5, a global average pooling layer, and a convolutional block containing a 1D convolution with 1 filter of size 256. The final output is a feature representation of the middle 200 bp region. The head maps this representation to a predicted scalar output for insulation, using five fully-connected layers, each with 64 neurons and a dropout of 0.2 for the first four layers.

### Hyperparameter tuning

To obtain the optimal model architecture and parameters, we tested for several hyperparameters: size of first convolutional filter *s*_1_, pooling type, number of intermediate convolutional blocks, and the number of convolutional filters in these blocks *s_si_*. From these initial experiments (data not shown), we observed that many of these parameters such as the number of convolutional filters in the intermediate blocks did not substantially impact model performance. The two key parameters that did affect model performance were the number of convolutional blocks *n_si_* and input size *w_c_*.

To verify whether model performance was saturated, we ablated combinations of these two parameters, training several models with *n_si_* being one of {1, 2, 3, 4, 5, 6, 7} and DNA context bin width *w_c_* being one of {8, 12, 16, 20}. Thereafter, we computed the validation performance between the dataset Domino was trained on and another dataset comprised of 8 independent Micro-C datasets in nc14 as a proxy for model saturation. As **Fig. S1** illustrates, model performance saturates for various models. Given that many models performed equally well (had the highest validation performance), we chose one such model with the hyperparameters *n_si_*= 5 and *w_c_* = 16.

### Domino training and evaluation

During model training, we used the Mean Squared Error (MSE) as our objective function, the AdamW optimizer with a learning rate of 10^−3^ and a batch size of 32. We used the ReduceLRonPlateau learning rate scheduler from PyTorch and early stopping with a patience of 20 using Pearson correlation as the monitored metric.

Following training, we evaluate model performance by computing the correlation between the true and predicted insulation values for the held-out test dataset. Additionally, we assessed model performance across entire regions using a “sliding window” approach. Specifically, Domino was used to predict the insulation score for every consecutive 200 bp bin in the region by sliding an input window by 200 bp in each step. This produced a predicted insulation track, to which the ground truth insulation track could be compared. For analysis in **Fig. S1B** and **Fig. S7C**, we considered insulation tracks in regions of size 30.2 Kb. Values than were within 400 bp of non-finite values in the ground truth insulation track were omitted from consideration when computing correlations between predicted and true insulation tracks. Additionally, all regions that yielded less than *n* = 10 values for comparison were removed.

### Predictions on mutagenized boundaries at the *eve* locus

We downloaded the Micro-C maps obtained from Ke et al. (2024)^30^. In the study, there were four separate experimental perturbations, in which the *nhomie* boundary (of approximately 600 bp in length) was replaced with one of the following:

1. *nhomie* forward (or reference): the endogeneous *nhomie* boundary was replaced with a copy of *nhomie* in the forward orientation.
2. *nhomie* inverted (or reference inverted): the endogeneous *nhomie* was replaced with a copy of *nhomie* in the reverse orientation (i.e., reverse complement).
3. *homie*: the endogeneous *nhomie* was replaced with a copy of the *homie* boundary.
4. lambda: the endogeneous *nhomie* was replaced with a 606 bp phage *λ* DNA fragment.

For each mutant, we first computed the insulation score using a window size of 1 Kb using the corresponding contact maps. Using customized genomes for all four mutants containing the experimental edits, Domino was used to predict the insulation score track centered on the *nhomie* boundary using a “sliding window” approach described earlier.

### Construction of the set of boundaries

First, we gathered a set of genomic sites with strong insulation, which consisted of all dataset examples that surpassed a boundary score threshold of 1.0; the boundary score is a readout of boundary strength from FAN-C. To determine this threshold, we used a strategy similar to that used in Ramírez et al. (2018)^14^ for analyzing binding levels of CP190, an architectural protein that was shown to bind to nearly all previously studied promoter and non-promoter boundaries in *Drosophila*^14^. First, we discretized all the 97,807 200 bp-long local minima sites called by our boundary calling procedure into boundary score intervals of width 0.1 (e.g., (0.0, 0.1], (0.1, 0.2] and so on). Next, CP190 ChIP-seq levels were quantified and Wilcoxon rank-sum tests were performed between all intervals. Because the interval of (0.9, 1.0] was the first interval that had a statistically significant difference in CP190 levels from lower intervals, we filtered for all examples that surpassed a boundary score cutoff of 1.0, which resulted in a preliminary set of 8, 002 strong boundaries. Thereafter, we removed all such boundaries where the balanced contact frequency in its bin and any bin within 800 bp of said boundary was not defined (which visually appears as a thin stripe in the contact map). In total, we obtained 7,311 boundaries.

For the larval and adult brain, we used a boundary score cutoff of 1.2 and 1.4 respectively; boundary set construction was otherwise the same. This yielded 8,443 and 8,164 larval and adult brain boundaries, respectively.

### *De novo* motif discovery using Domino sequence attributions

We used the *in silico* mutagenesis (ISM) tool from the tangermeme^33^ package to compute attribution scores for boundary sequences. To systematically uncover insulator motifs from Domino attribution scores, we used the tfmodisco-lite^76^ package implementation of the TF-MoDISco^31^ algorithm. Using the attribution scores computed for every base-pair, TF-MoDISco was used to identify “seqlets”, or spans of nucleotides with high attribution magnitudes before clustering them into motif representations called contribution weight matrices (CWMs).

Because the majority of the sequence attribution were in the middle 400 bp of boundary sequences (**Fig. 4B**), we used the attributions in the center 600 bp of embryonic boundary sequences as input to TF-MoDISco. In total, 63 preliminary seqlet clusters (or hereafter referred to as “motifs”) were discovered, 27 with positive attributions (weaken predicted insulation) and 36 with with negative attributions (strengthen predicted insulation); for notation purposes, we denote the former and latter as positive motifs and negative motifs respectively. All negative motif patterns were filtered to have at least 85 seqlet instances; all positive motif patterns were filtered to have at least 100 seqlet instances and a minimum average attribution of 0.02. Additionally, motifs with a total information content (across all positions) of less than 7 were discarded. After filtering, we obtained a total of 24 negative motifs and 2 positive motifs. For motif discovery in the larval brain, we imposed an average attribution cutoff of 0.015 for positive motifs. For the adult brain, we imposed an average attribution cutoff of −0.03 and 0.025 for negative and positive motifs, respectively. A cutoff of 100 seqlets were also used for all motifs in both the larval and adult brain. In total, 26 negative and 5 positive motifs were identified in the larval brain; 34 negative and 13 positive motifs were identified in the adult brain.

To characterize which binding factors matched our discovered Domino motifs, we used several motif sources. First, Domino-derived motifs were matched against all insect motifs from the 10^th^ release (2024) of JASPAR^77^ using the tfmodisco-lite tool^76^. Then, Domino-derived motifs without significant matches were further matched against motif predictions (PWMs) for a collection of zinc-finger arrays from zf.princeton.edu^50,51^. Significance for these matches was assessed using the tomtom-lite package^78^ implementation of the TOMTOM algorithm^79^. For matching of several *Lola* isoforms discovered in the brain and for additional validation of other factor candidates, we obtained protein sequences for C2H2 factor candidates from UniProt (uniprot.org)^80^ and generated the corresponding PWM predictions from zf.princeton.edu^50,51^. Finally, for motifs that were not present in any of the previously described sources, we manually generated the PWM representation of motifs that were published in prior literature with their respective sources shown in **Fig. 2C**.

For analysis involving finding motif occurrences within boundary examples, we used the FIMO implementation from tangermeme^33^ using the default parameters.

### Processing of ChIP-seq data

We downloaded raw ChIP-seq data from the Sequence Read Archive (SRA) database FASTQ (**Table S1**). Reads were first trimmed using Trim Galore^81^ and then aligned to the reference genome with BWA-MEM^67^ using the default parameters. Then, SAMtools^82^ was used to sort the BAM alignments followed by filtering out supplementary, non-primary, and unaligned reads. Then, Picard Tools^83^ was used to filter out duplicate reads. Finally, BAM alignments for replicates of the same condition were merged using SAMtools and merged bam files using bamCoverage tool from deepTools^84^.

### Clustering of motif attributions

To cluster motif attributions across our boundaries, we first computed attribution scores for all the discovered motif instances among all 7, 311 boundaries. Specifically, for each of the 24 primary motifs, attribution of the motif instance (or instances if there was more than one) present in the middle 600 bp of every boundary example was summed to obtain a total attribution for the motif. To account for both the positive and negative strands, the average of the total motif attributions of each strand was computed to obtain an attribution matrix, with 7, 311 boundaries (rows) and and 24 motifs (columns). We then filtered for all boundaries with nonzero attributions for the negative primary motifs, resulting in a final attribution matrix of 4, 766 boundaries. Among these, a total of 4, 289 boundaries had at least one motif with sufficiently strong total attribution (*<* −0.25) for at least one primary motif; unless otherwise specified, we used this matrix *M* (4,289 boundaries by 24 motifs) for clustering analysis (results presented in **Fig. 5**).

Next, we computed a binary matrix *B* on *M*, where an element *B_i j_*was 1 if total attribution for motif *j* in boundary *i* was less than −0.25; otherwise, *B_i j_* was 0. Standardization and K-means clustering was performed on *B* using *K* = 30 to generate cluster assignments for all boundaries; the parameter *K* = 30 was chosen based on systematic testing of *K* ranging from 2 to 75 (**Fig. S5A**). The cluster assignments were then used to order the rows for the final sorted motif clustermap.

For each cluster, we performed several analyses. To compute the fraction of boundaries that overlapped with transcriptional start sites (TSS), we downloaded the reference gene annotation file (dm6.refGene.gtf.gz) using the dm6 genome build from the UCSC Genome Browser^85^. After filtering for all transcripts that corresponded to a mRNA transcripts for protein coding genes, the position of the TSS was identified based on TSS direction. Finally, the fraction of boundaries that was within 1 Kb of TSS for each cluster was calculated.

We repeated the same procedure to cluster motif attributions from Domino models trained on Micro-C data from the larval and adult brain. For the larval brain, clustering was performed on boundaries across 26 primary motifs using *K* = 30; for the adult brain, clustering was performed on boundaries across 34 primary motifs using *K* = 40.

### Statistical analysis of general and pairwise motif co-activity

For general motif co-activity, we examined statistical enrichments for a motif being co-active with other motif. A two-sided Fisher’s exact test was performed using the set of all boundaries with some negative primary motif attribution (*n* = 4, 766) across all 24 primary motifs according to the following contingency table:

|  |  |
| --- | --- |
| Active motif X with another active motif | Active motif X without another active motif |
| No active motif X, another active motif | No active motif X nor another active motif |

For pairwise motif co-activity enrichment, we then examined enrichments for a specific pair of motifs being co-active. A two-sided Fisher’s exact test was performed using the same boundaries and motifs used for general motif co-activity analysis. The following contingency table was used:

|  |  |
| --- | --- |
| Strong motif X and motif Y | Strong motif X, no strong motif Y |
| No strong motif X, strong motif Y | No strong motif X nor motif Y |

For pairwise motif co-activity analysis in the *Drosophila* brain, we used the same procedure described above using the set of boundaries shown in their respective clustermaps (**Fig. S10**).

For all co-activity analyses, adjusted *p*-values were computed using the Benjamini-Hochberg correction procedure.

### *In silico* motif synergy analysis

For motif cooperativity, we defined synergy score using a formulation similar to the approach of the previous published model DeepSTARR^54^. First, we randomly sampled 100 “backbone” sequences of length 6,600 bp from the *Drosophila* genome among examples that were not local minima in our dataset. Then, among all primary motifs, we took the 20 motifs with the most frequent negative attribution (**Fig. S5D**) and computed the synergy score between all 400 possible ordered motif pairs (including a motif with itself).

For an ordered motif pair (*A, B*), we represented *A* and *B* by their corresponding consensus sequences. Specifically, we inserted motif *A* into the center of a backbone and *B* at varying distances downstream from the central motif. We predicted insulation for just the backbone (*y_b_*), the backbone with motif *A* (*y_A_*), the backbone with motif *B* without *A* (*y_B_*), and the backbone with *A* and *B* (*y_AB_*). The synergistic effect of *B* on *A* is defined as the relative change between the predicted insulation of having *A* and *B* both present and the sum of their predicted insulations separately. Mathematically, this can be represented in the following way:

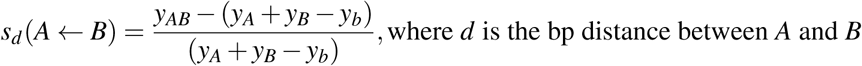

We define synergy as a relative change since it would provide the ability to directly compare synergy scores between different ordered pairs of motifs.

We computed the synergy scores between all ordered pairs of motifs at distances ranging from 1 to 150. For a given distance, we defined the synergy score *S_d_*(*A* ← *B*) of *A* ← *B* at distance *d* to be the median of the 100 synergy scores *s_d_* across all the backbones. Finally, to obtain an aggregate level of synergy for *A* ← *B*, we defined the final synergy level for *A* ← *B* to be the mean of the 20 largest synergy scores.

### CG4854, Trem, and CG17385 peak calling and *de novo* motif discovery

We downloaded ChIP-seq data for CG4854, Trem, and CG17385 from ENCODE as FASTQ files (**Table S1**). Each of the three factors had two biological replicates and one input control used for peak calling. For each factor, we used Bowtie2^86^ to align all reads to the *Drosophila melanogaster* genome assembly version 6 (dm6) reference genome using the –very-sensitive flag. Next, we used SAMtools^82^ to sort, de-duplicate and index the alignments. We then used MACS2^87^ to call peaks using the respective input control, an effective genome size of 142, 573, 017, and a q-value threshold of 0.05, and Signal Per Million Reads option with no additional filtering of duplicates. Summits within peaks were called. Finally, using peaks called at a q-value threshold of 0.05, we used the IDR package (https://github.com/nboley/idr)^88^ to filter for reproducible peaks between the two replicates using a global IDR threshold of 0.05. In total, this yielded 2,581 and 1,044 reproducible peaks for Trem and CG4854, respectively.

We performed *de novo* motif discovery within these peaks using MEME-ChIP^53^. Specifically, we first used the getfasta tool from BEDTools^89^ to extract the 400 bp-long sequences centered on the summits within the 1000 most significant IDR thresholded peaks. Then, we ran MEME-ChIP using a maximum dataset size to be 3 × 10^6^ and a maximum sequence length of 200 bp (i.e., consider only the center 200 bp of each sequence).

### CG4854 and Trem peak and boundary analysis

For peak set analysis, we used the intersect tool from BEDTools^89^ using the pybedtools package^90^ for CG4854 and Trem peaks. We considered two peaks (500 bp) to overlap with one another if one peak overlapped at least one base pair of the other peak. Similarly, a peak overlapped a boundary (200 bp) if they overlapped in at least one base pair. For assigning insulation score to a peak, if the peak overlapped a boundary, insulation of the boundary was assigned to the peak; otherwise, if a peak did not overlap a boundary but overlapped multiple bins, the strongest (most negative) insulation value was assigned to the peak.

To test whether Trem and CG4854 peaks were enriched in boundaries, we performed a permutation test on the peaks. For both factors, locations of peaks were randomly permuted to locations in the genome which did not overlap with blacklisted regions^73^. The proportion of permuted peaks that overlapped boundaries was then computed. This procedure was repeated 10,000 times to form an empirical null distribution. The empirical p-value was the fraction of permutations where the permuted peak-boundary overlap was greater than the observed peak-boundary overlap.

### HCR-FISH

The sequences of target genes were obtained from Flybase (flybase.org)^91^. To design probes, the target gene sequences were submitted to the Molecular Instruments probe design platform (www.molecularinstruments.com/hcr-rnafish)^92^, with parameters set to a 35 probe set size for *Drosophila melanogaster*. A similar method was designed based on published smFISH methods^93,94^. 100-200 flies were placed in a cage with an apple juice plate at the bottom of the cage. For st14 embryos, collections were done overnight. Embryos from each plate were washed into collection mesh and dechorionated in bleach for 2 min, then fixed in 5 mL of 4% paraformaldehyde in 1X PBS and 5 mL of heptane for 15 min with horizontal shaking. The paraformaldehyde was then removed and replaced with 5 mL methanol. The embryos were then devitellinized by vortexing for 30 s, and washed in 1 mL of methanol twice. Methanol was then removed and replaced by PTw (1X PBS with 0.1% Tween-20) through serial dilutions of 7:3, 1:1, and 3:7 methanol:PTw. The embryos were washed twice in 1 mL of PTw and pre-hybridized in 200*µ*L of probe hybridization buffer for 30 min at 37°C. 0.4 pmol of each probe set were added to the embryos in probe hybridization buffer, and the embryos were incubated at 37oC for 12-14 h. The embryos were then washed 3X with probe wash buffer at 37°C for 30 min and 2X with 5X SSCT(5X SSC+0.1% tween) at room temperature for 5 min. Then the embryos were pre-amplified with 300 *µ*L amplification buffer for 10 min at 25°C. Meanwhile, 6 pmol of hairpin h1 and h2 were snap-cooled separately (95oC for 90 s, cool to RT with a 0.1oC drop per second), and then mixed in 100 *µ*L of amplification buffer at room temperature. After that, the pre-amplification solution was removed from the embryos, and 100 *µ*L of hairpin h1/h2 mix were added to the embryos. Next, the embryos were incubated for 12-14 h at room temperature in the dark. To remove excess hairpins, the embryos were washed in SSCT as follows: 2X for 5 min, 2X for 30 min, and 5X for 5 min. Then, the embryos were washed with 1 mL PTw for 2 min and stained with DAPI/Hoechst at 1 *µ*g/mL for 15 min at room temperature in the dark. The embryos were then washed with PTw 3X for 5 min. Finally, the embryos were mounted on microscope slides with Vectashield and a #1.5 coverslip for imaging.

### Imaging and image analysis

Embryos from smFISH were imaged using a Nikon A1 confocal microscope system with a Plan Apo 20X/0.75 DIC objective. Z-stack images were taken at intervals of 2 *µ*m, 4X average, 1024×1024 resolution, and the appropriate laser power and gain were set for the 405, 561, and 640 channels to avoid overexposure. Images were processed using ImageJ, and the maximum projection was applied to each of the stack images. For APR intensity measurements, the ROI tool was used to crop out the APR region of late-stage embryos. The cells with APR signal were also highlighted and selected by MaxEntropy thresholding. The particle measurement tool was used to measure the average intensity of all cells that had a signal. At the same time, the background signal (average intensity) was taken from cells without a signal in the same embryo. The relative intensity (signal to background) for each embryo was calculated using the stripe signal and background from the same embryo. To make comparisons between independent biological replicates, the average background signal of all embryos from each replicate was calculated. The relative intensity of each embryo from each replicate was normalized based on the average background signal of all embryos from that replicate. GraphPad Prism was used for data visualization and statistical analysis. To compare intensity from embryos in different groups, different signals from the same embryo (e.g., *LacZ* and *GFP*) were paired, and paired two-tailed *t*-tests were used to calculate p-values. Non-parametric unpaired *t*-tests were used to calculate percent embryos that show *LacZ* signal in the *hebe* activated cells.

## Supplementary information

### Supplementary figures

**Figure S1.**
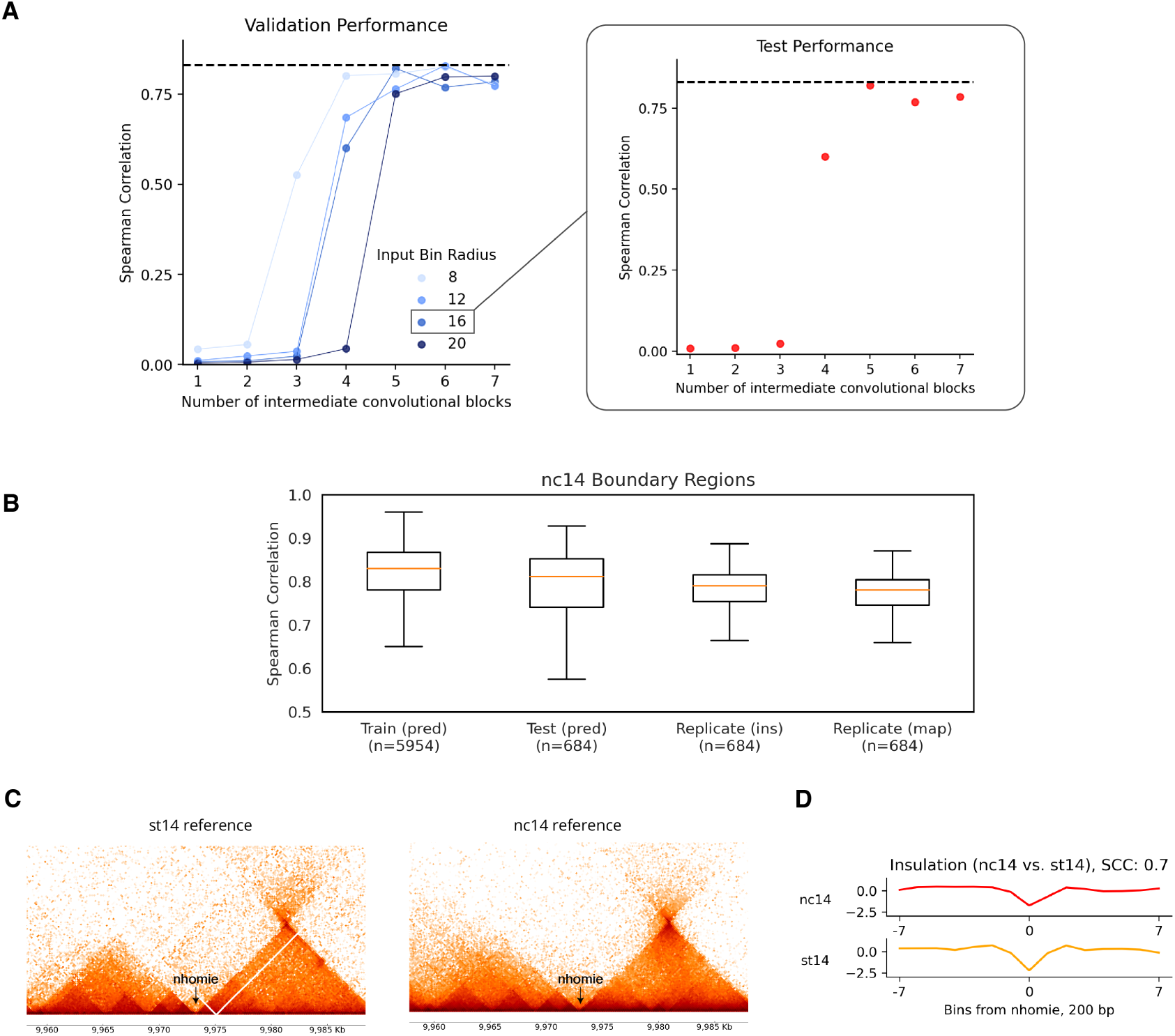
Optimization and evaluation of the Domino framework. (**A**) Hyperparameter sweep exploring the number of convolutional layers and sequence context length, evaluated on the validation set (left) and held-out test set (right). The input bin radius denotes the number of 200 bp bins flanking the central 200 bp target bin. The inset shows the test performance across different values for the number of convolutional blocks for a fixed input bin radius = 16 that showed the highest validation performance. The dashed line represents the correlation observed between independent biological replicates. (**B**) Evaluation of predicted insulation tracks across 30.2 Kb genomic regions. Boxplots show the distributions of Spearman correlations between predicted and observed insulation scores for 30.2 Kb regions centered on boundaries from the train set and test set, as well as correlations between biological replicates (aggregated 11 datasets used in training vs. aggregated independent 8 datasets, as in **Fig. 1C**). Also shown is the distribution of correlations between 2D Micro-C contact maps at 200 bp resolution across the same test regions. Boxplots: center line, median; box limits, upper and lower quartiles; whiskers, 1.5× interquartile range; outliers not shown. (**C**) Micro-C contact maps at the *eve* locus compared between the st14 and nc14 embryonic stages, centered on the *nhomie* boundary. (**D**) Comparison of insulation scores within the 1.6 Kb region centered on the *nhomie* boundary between nc14 (top) and st14 (bottom) embryonic stages. SCC, Spearman corelation coefficient.

**Figure S2.**
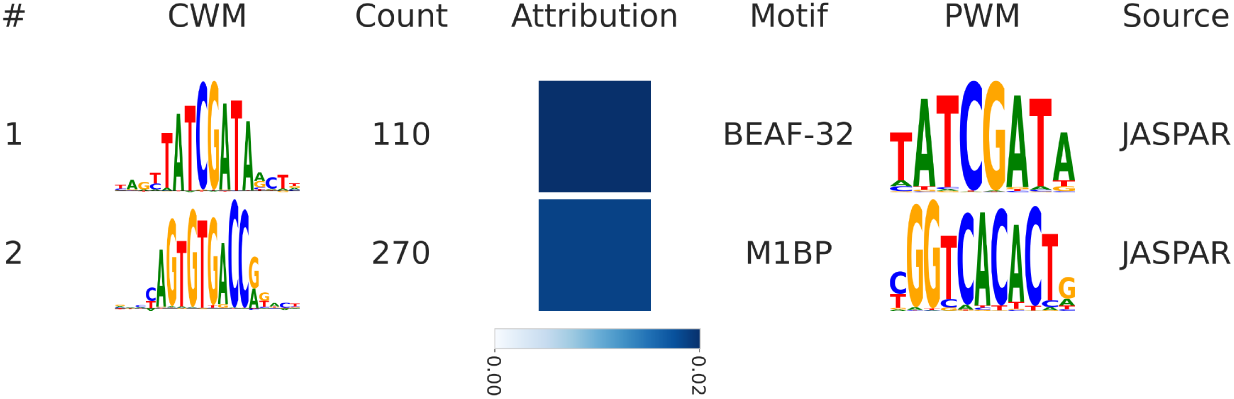
Motifs with positive attributions discovered *de novo* by Domino in the nc14 embryo. Motifs are ranked by average attribution. Each row displays a motif identifier, CWM from TF-MoDISco, seqlet count, average attribution score, and the best match to known motifs (motif name, PWM logo, and source of the motif).

**Figure S3.**
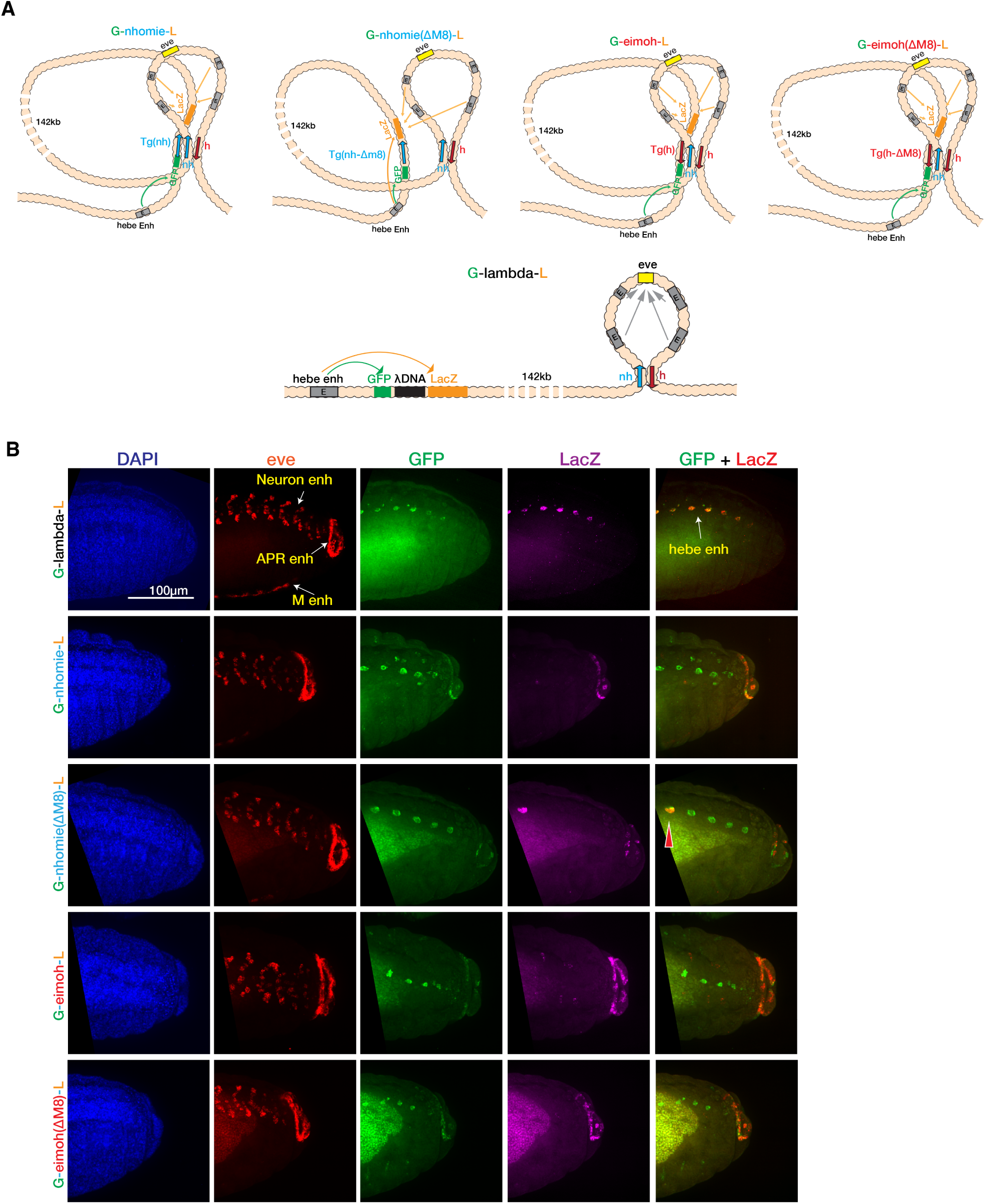
Proposed chromosomal topologies for *hebe* transgene reporter assays. **(A)** Schematic representations of the proposed topologies for each of the five integrated constructs. For the *nhomie* and *homie* models, the transgene element pairs with endogenous *nhomie* and *homie* in a configuration where the *hebe* enhancer preferentially interacts with *GFP* while *eve* enhancers preferentially interact with *LacZ*. For the *homie(*Δ*M8)* model, the configuration is similar given that Motif-8 is not active in *homie*. In contrast, for the *nhomie(*Δ*M8)* model, deletion of Motif-8 weakens pairing and insulation, resulting in reduced interactions between *eve* enhancers and *LacZ* but strengthened interactions between *hebe* enhancer and both *LacZ* and *GFP*. For the *lambda* control, pairing does not occur between the transgene element and the *eve* locus and insulation is absent; consequently, the *hebe* enhancer activates *LacZ* and *GFP*, while independently the *eve* enhancers activate *eve*. **(B)** Representative examples of HCR-FISH embryo microscopy images arranged in columns (left to right): DAPI, *eve*, *GFP*, *LacZ* and *GFP* + *LacZ* expression, across the transgene strains (rows). Yellow arrows mark the locations of active enhancers. The red arrow indicates the specific embryonic region with *hebe* enhancer-activated *LacZ* expression. APR, anal plate ring; M, mesoderm.

**Figure S4.**
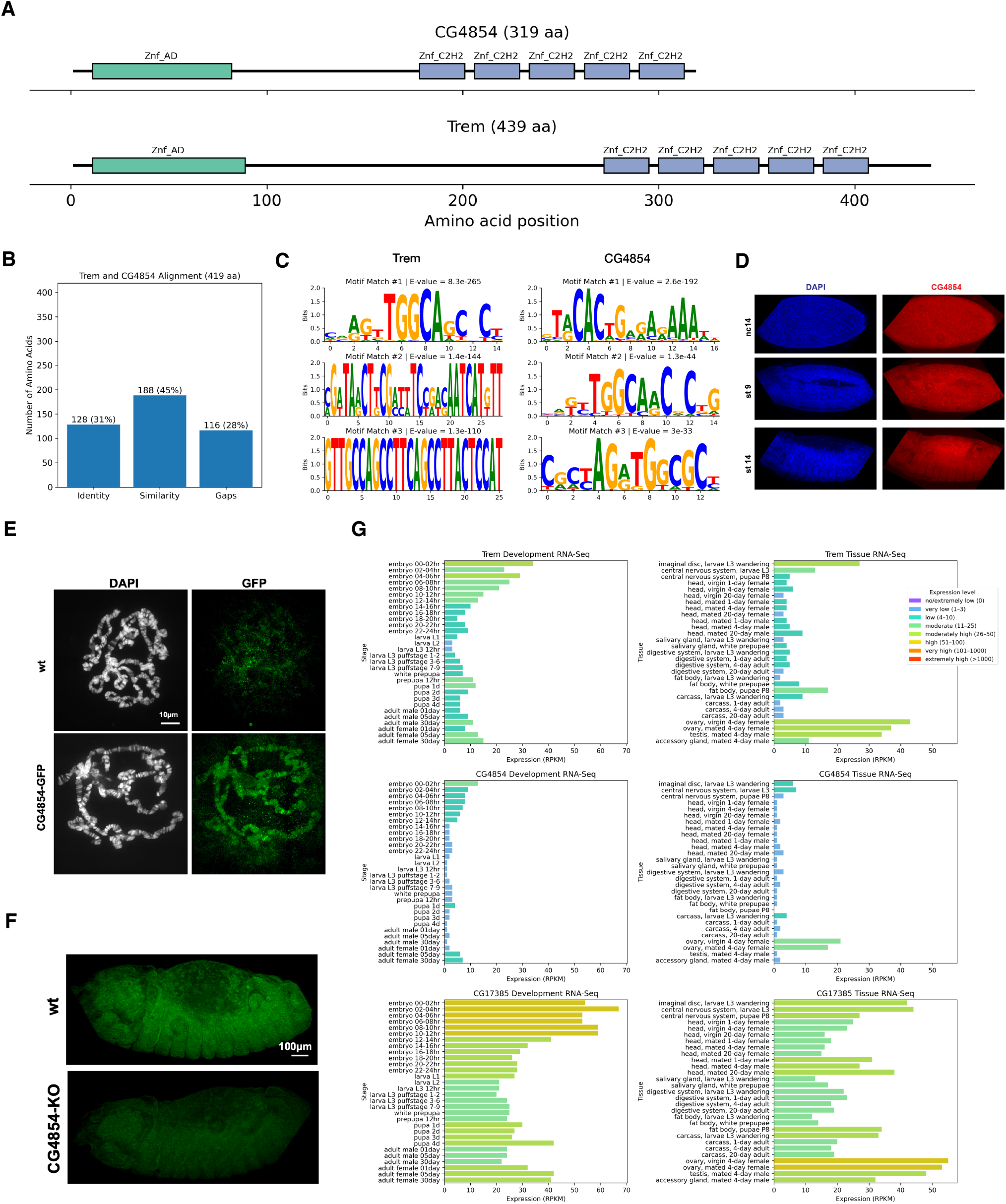
Characterization of CG4854, Trem and CG17385. (**A**) Domain architecture of CG4854-PA (319 amino acids) and Trem-PA (439 amino acids) protein isoforms from the SMART database^95^ obtained from FlyBase^96^ (release FB2025_04). Znf_AD, Zinc Finger-Associated Domain; Znf_C2H2, Zinc Finger Cys_2_His_2_ domain. **(B)** Sequence alignment statistics comparing CG4854 and Trem protein sequences. Percentages are calculated relative to the total alignment length of 419 amino acids. Identity and similarity indicate the number of exact amino acid matches and the number of biochemically or structurally similar matches, respectively. Statistics were obtained from the DIOPT Ortholog Finder Tool (Version 10.0)^97^. **(C)** Motifs discovered by MEME-ChIP within the Trem (left) and CG4854 (right) ChIP-seq peaks, ranked by significance of E-value. The Domino-derived motif #6 in the list of motifs with negative attribution (that matches previously described Motif-8^14^) resembles the Trem motif match #1 and CG4854 motif match #2. **(D)** Ubiquitous embryonic expression of CG4854 throughout embryonic development. HCR-FISH visualization of CG4854 transcripts in wild-type (WT) embryos across nuclear cycle 14 (nc14), stage 9 (st9), and stage 14 (st14) (top, middle, and bottom rows, respectively). Blue, DAPI; red, CG4854 transcripts. **(E)** Immunostaining for GFP on salivary gland polytene chromosomes from WT and transgenic larvae expressing a CG4854-GFP fusion protein (y1 *w*^∗^; PBac{CG4854-GFP.FPTB}VK00037) at the third instar stage. CG4854-GFP localizes to chromatin. **(F)** HCR-FISH detection of CG4854 transcripts in wild-type (WT) and *CG4854* knockout (*w*^1118^; PBacWHCG4854f02794) stage 14 embryos. Only low-to-background levels of CG4854 signal were detected in *CG4854* KO embryos. **(G)** Expression patterns of Trem (first row), CG4854 (second row), and CG17385 (third row) across developmental stages (first column) and across tissues (second column). Processed RNA-seq gene expression data was obtained from FlyBase (flybase.org)^91^.

**Figure S5.**
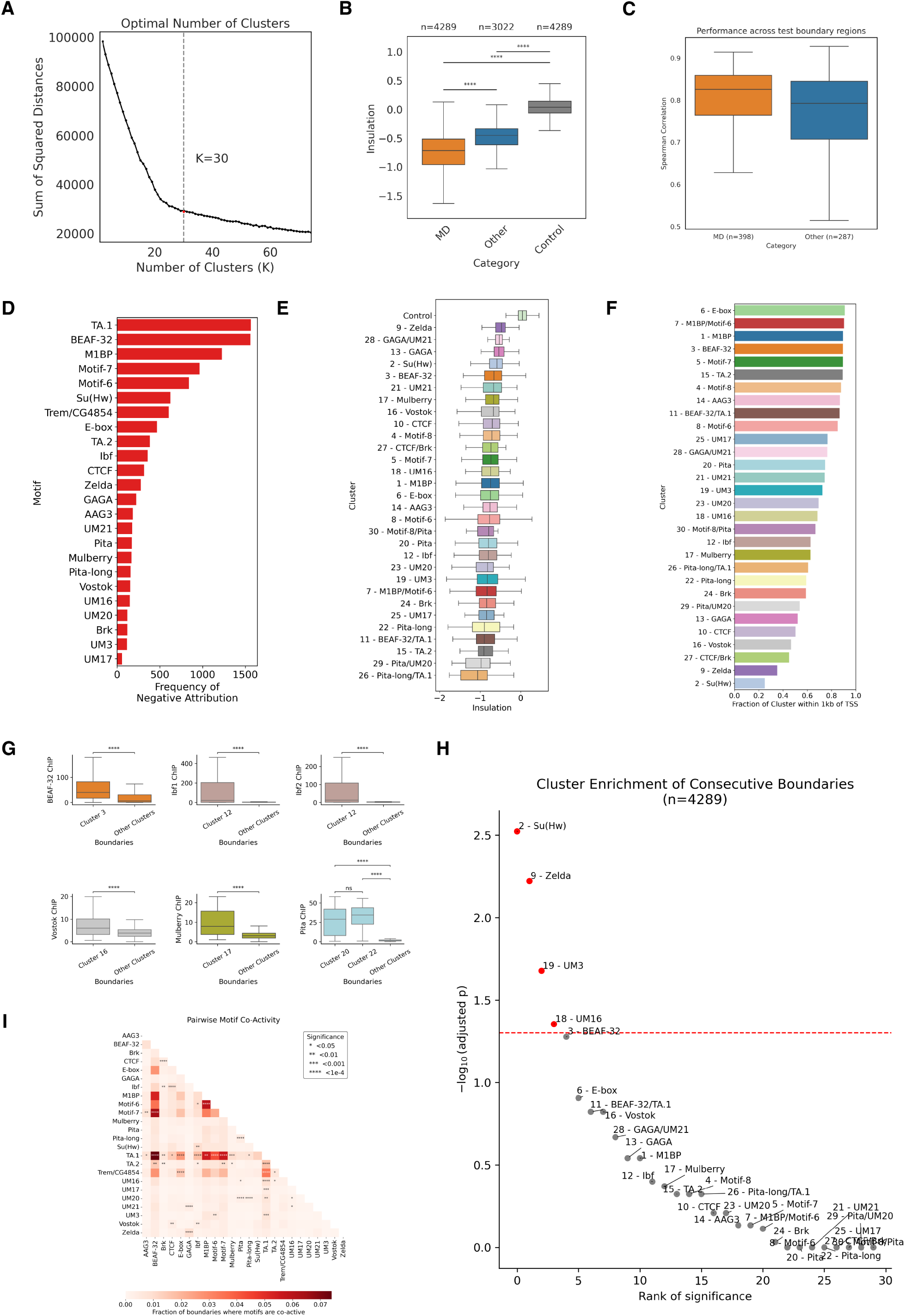
Characterization of boundary motif clusters. (**A**) Elbow plot of *K*-means clustering for the number of distinct boundary clusters *K* ranging from 2 to 75. The red dashed line marks *K* = 30, the number selected for downstream analysis. (**B**) Distribution of insulation scores across all boundaries, comparing those that have an active negative attribution motif (motif-dependent, MD) against all other boundaries and control regions (*n* = 4, 289 random 200 bp non-boundary genomic sites). Significance is assessed using a one-sided Wilcoxon rank-sum test; ****, *p <* 0.0001. (**C**) Evaluation of predicted insulation tracks across held-out test boundary 30.2 Kb regions (as in **Fig. S1D**), stratified by boundary type (motif-dependent vs. other). (**D**) Number of boundaries with negative attribution for all 24 negative attribution motifs. (**E**) Distribution of insulation scores across all clusters. Control regions include *n* = 4, 289 random 200 bp non-boundary genomic sites. (**F**) Fraction of boundaries in each cluster that are within 1 Kb of a transcription start site (TSS) of a protein coding gene. (**G**) ChIP-seq signal enrichment at selected clusters. Comparison of insulator factor ChIP-seq signal intensity between boundaries in the cluster corresponding to the factor and boundaries from all other clusters. Significance is assessed using a one-sided Wilcoxon rank-sum test; ****, *p <* 0.0001. (**H**) Statistical enrichment of co-occurrence of boundaries from the same cluster as adjacent motif-dependent boundaries (*n* = 4, 289) in the genome. For each cluster, the number of adjacent boundary pairs belonging to this cluster were computed. Significance of this frequency was tested using a null distribution where all cluster labels (within the same chromosome) were randomly shuffled 10,000 times. The *p*-value was computed as the proportion of permuted number of occurrences that surpassed the observed number of occurrences. Benjamini-Hochberg correction was used to compute the adjusted *p*-value, or false-discovery rate (FDR). The red dashed line indicates a significance threshold of FDR = 0.05. (I) Comprehensive pairwise motif co-activity analysis. Fraction of all boundaries with negative primary motif attribution (*n* = 4, 766) where both motifs are active (strong attribution). *p*-values are obtained from two-sided Fisher’s exact test. Boxplots in this figure: center line, median; box limits, upper and lower quartiles; whiskers, 1.5× interquartile range; outliers not shown.

**Figure S6.**
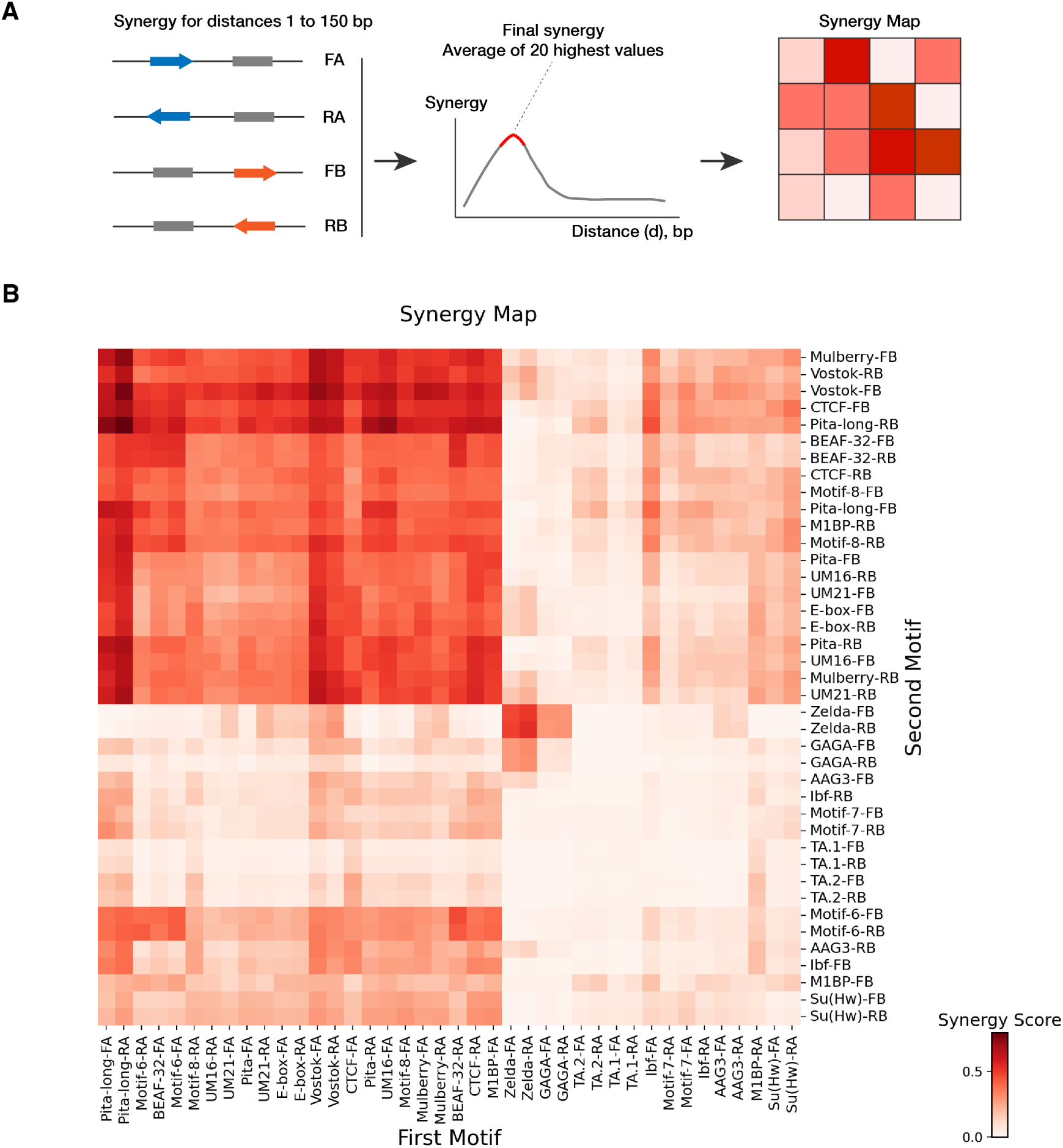
Motif synergy analysis. (**A**) Calculation of the motif synergy map, which summarizes synergistic relationships between pairs of motifs. For each motif pair, the synergy value is defined as the average of the 20 highest synergy scores across various distances. See **Fig. 6** and **Methods** for details. **(B)** Motif synergy map for all pairs of top 20 negative attribution motifs with highest attribution frequency. Motif positioned first are shown in columns, motifs positioned second (downstream of the first motif) are shown in rows.

**Figure S7.**
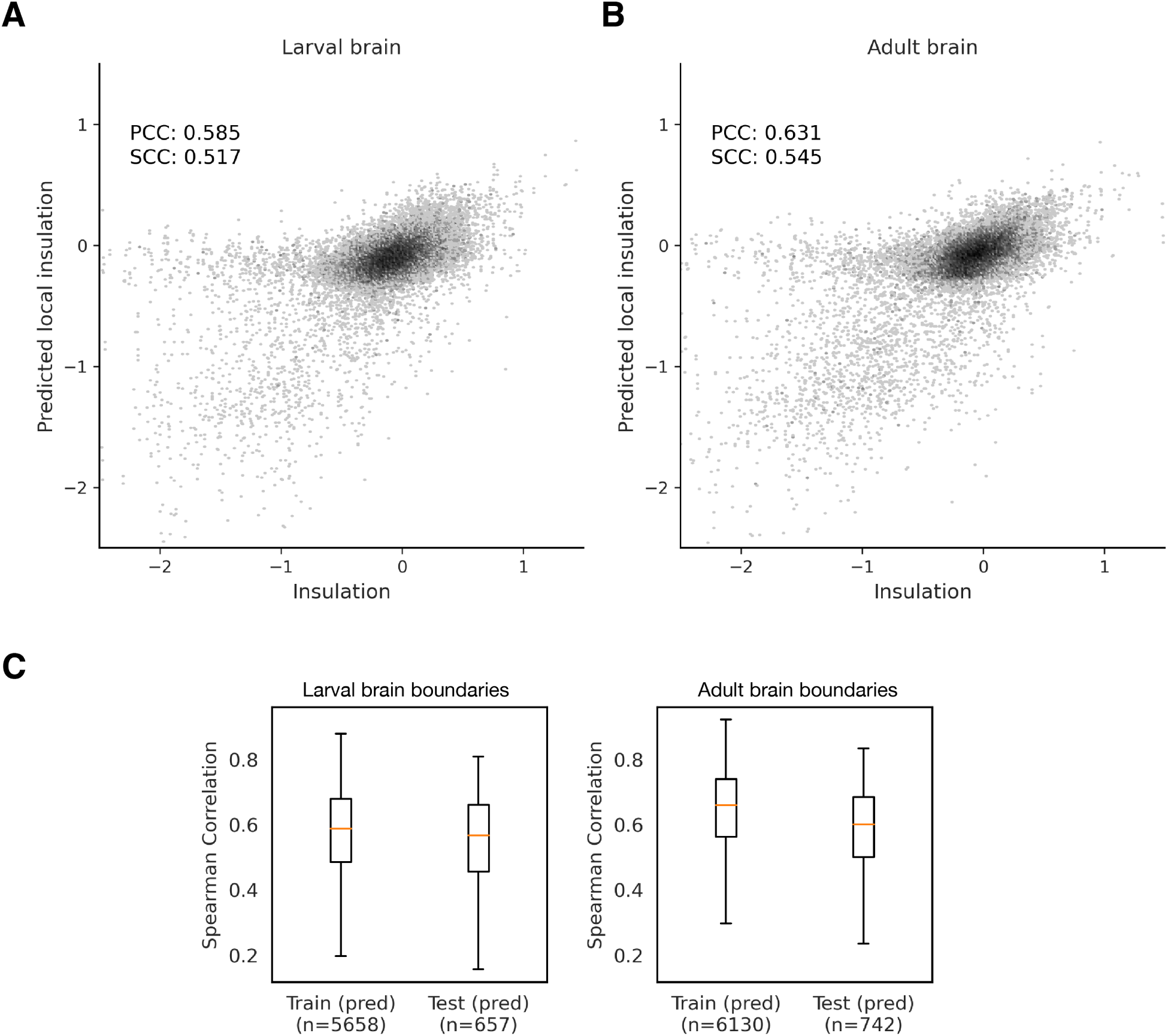
Domino performance for the larval and adult brain. (**A–B**) Model performance on held-out test data in the larval brain (A) and adult brain (B). Scatterplots show comparison between predicted and observed insulation. For visualization, outliers representing 0.21% of data points for larval brain and 0.20% of data points for adult brain are not shown. **(C)** Evaluation of predicted insulation tracks across 30.2 Kb genomic regions for larval and adult brain data (as in **Fig. S1B**).

**Figure S8.**
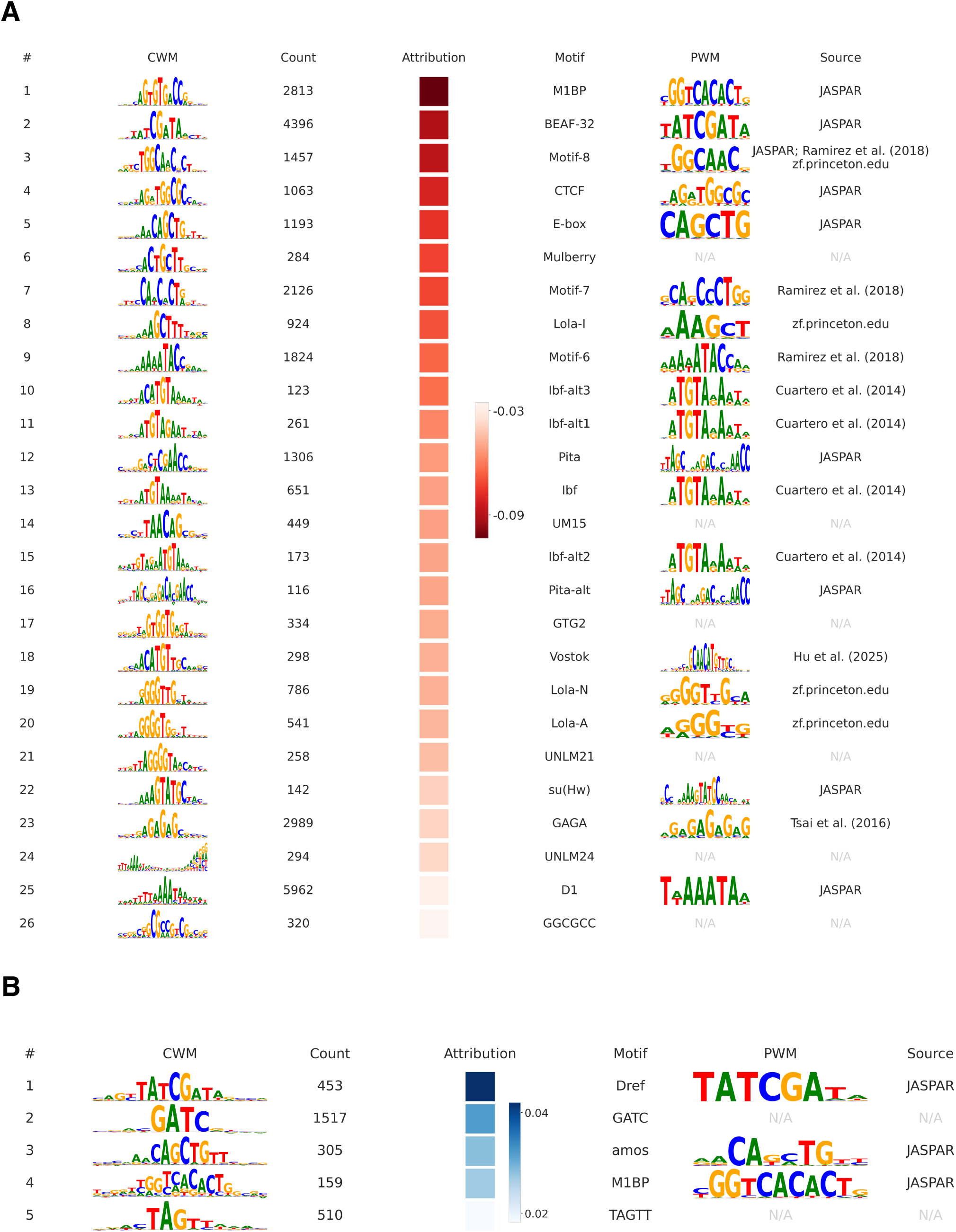
Motifs discovered by Domino in the larval brain. (**A–B**) Motifs discovered *de novo* by Domino from the larval brain boundaries. Motifs with negative attribution (A) and positive attribution (B), ranked within each category by average attribution. Each row displays a motif identifier, CWM from TF-MoDISco, seqlet count, average attribution score, and the best match to known motifs (motif name, PWM logo, and source of the motif).

**Figure S9.**
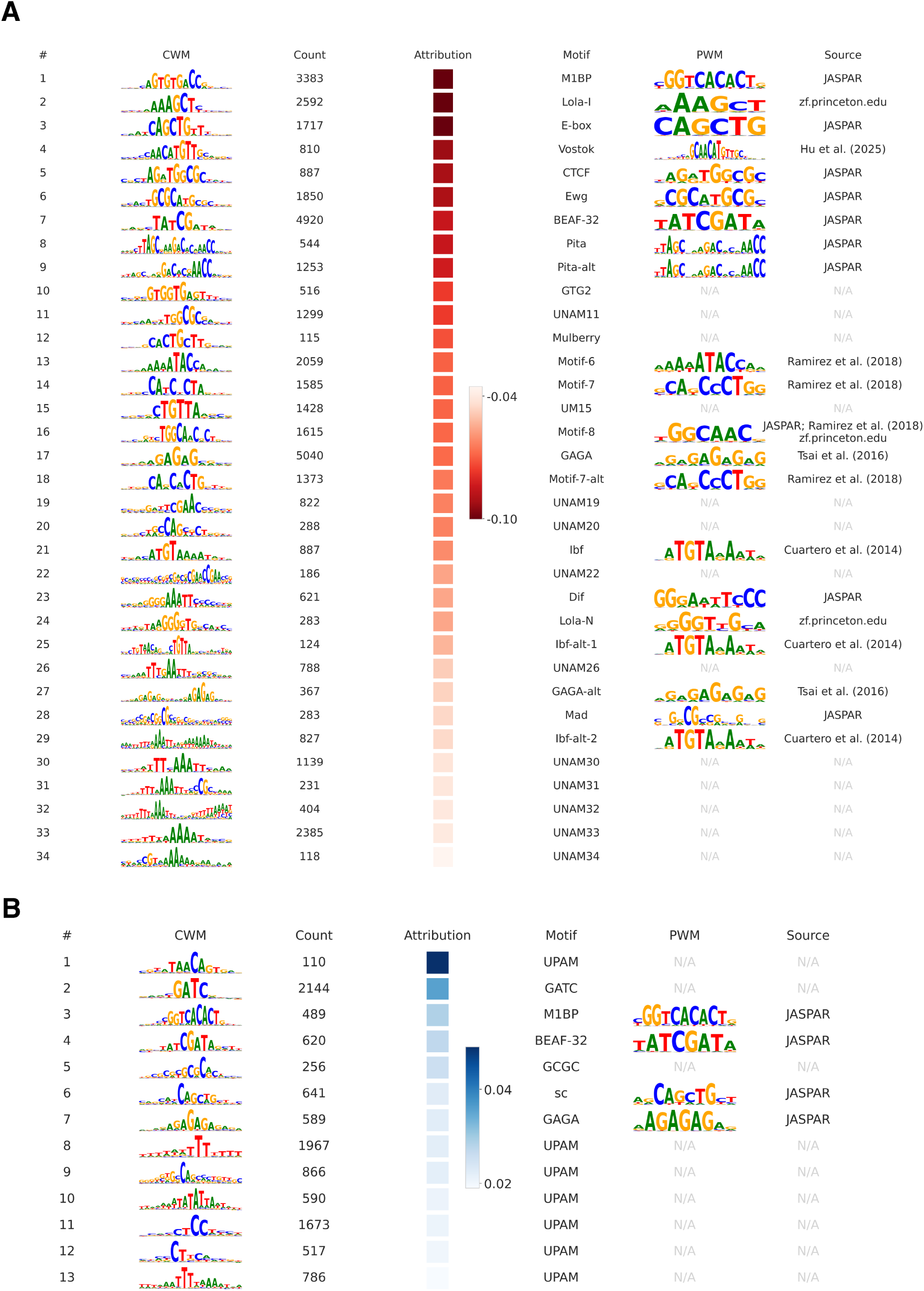
Motifs discovered by Domino in the adult brain. (**A–B**) Motifs discovered *de novo* by Domino from the adult brain boundaries. Motifs with negative attribution (A) and positive attribution (B), ranked within each category by average attribution. Each row displays a motif identifier, CWM from TF-MoDISco, seqlet count, average attribution score, and the best match to known motifs (motif name, PWM logo, and source of the motif).

**Figure S10.**
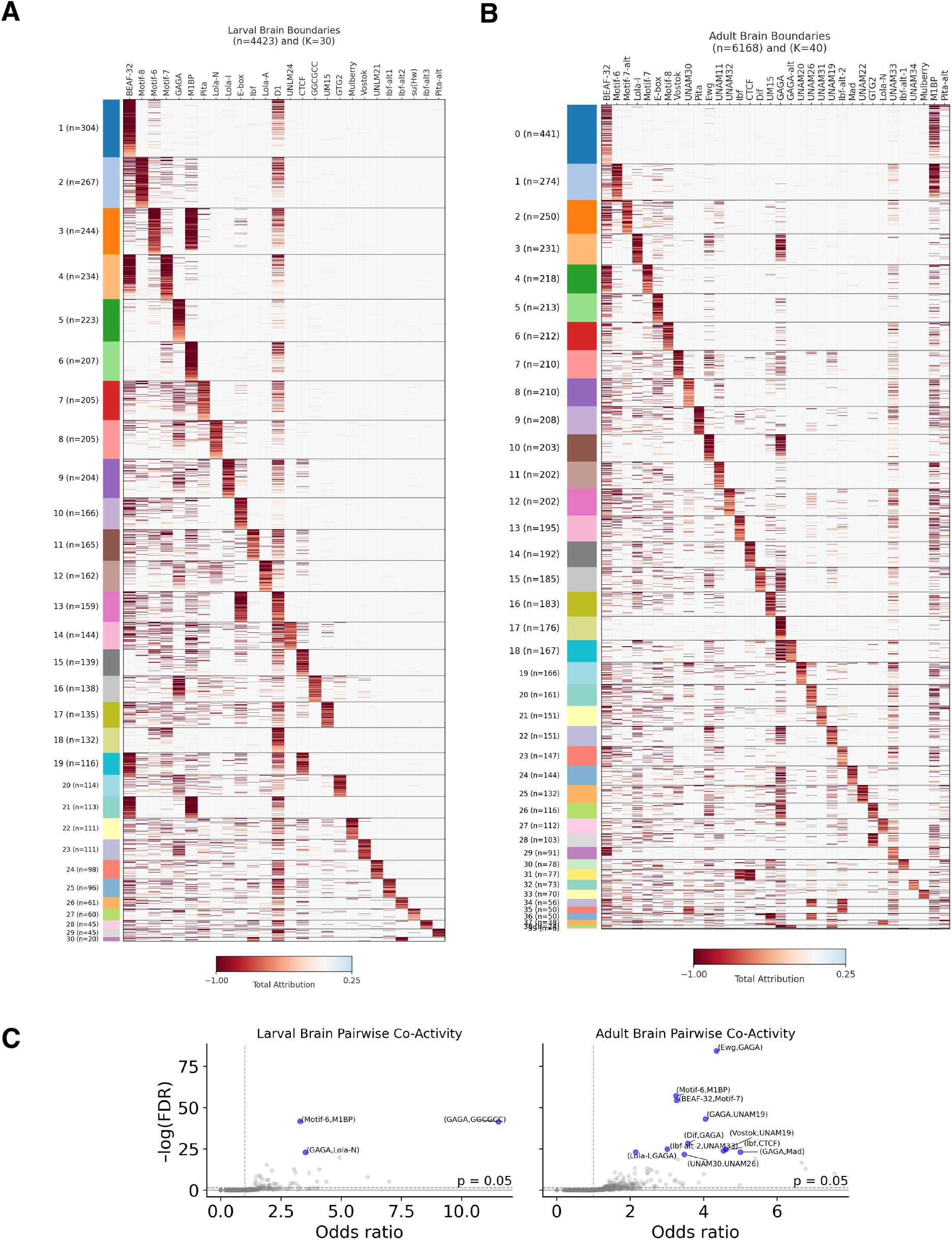
Tissue-specific insulation motif repertoire. (**A–B**) Clustermaps of the insulation motif repertoire for the larval brain (A) and adult brain (B). Same analysis as in **Fig. 5A**. **(B)** Pairwise co-activity between all pairs of motifs within strong larval brain-specific (left) and adult brain-specific (right) boundaries. Significance at FDR = 0.05 (horizontal dashed line) was assessed using a Fisher’s exact test.

### Supplementary table captions

Table S1: Micro-C and ChIP-seq data used in this study.

### Supplementary files

File S1: DNA sequences of transgene constructs with wild-type and Motif-8 variations of *nhomie* and *homie*.

File S2: Motif predictions for C2H2 4-ZF arrays.

