## Supplementary File S1 for "Sequence-to-function modeling uncovers the context-specific grammar of *Drosophila* chromatin insulation"

**homie WT**

AATACTAAAAAGTTTTTACTAGCATAAGCTGCGATTGAGCAGTTATTGTAGGGACAGGTTGATGGC
CGAGTGTTCTAGGGAAATGAATGAATGAAGATTTTTTCTTACTACCAAAACTTCTACAACTAAGTGA
AATAGACGGATTGAGTTCTACATACTTCCTACATATATTTTAGATATAGACATAGCCGAAAAGTATGC
TGCACTTTTTCAATAAAAACGTAAGCAGCTAAGCAGCGTAAGGCAGCTTCATAGCAGCGATCATC
TGCCAGCGAGCATAGCCAAGTTAGGCACCTGCACACATTGCCACTCTTTCTTTTAGCGTTGCCA
CTTCAATTTCTTTTACAAACCATCGCAGCGTGTAAT

**homie Δmotif-8**

AATACTAAAAAGTTTTTACTAGCATAAGCTGCGATTGAGCAGTTATTGTAGGGACAGGTTGATGGC
CGAGTGTTCTAGGGAAATGAATGAATGAAGATTTTTTCTTACTACCAAAACTTCTACAACTAAGTGA
AATAGACGGATTGAGTTCTACATACTTCCTACATATATTTTAGATATAGACATAGCCGAAAAGTATGC
TGCACTTTTTCAATAAAAACGTAAGCAGCTAAGCAGCGTAAGGCAGCTTCATAGCAGCGATCATC
TGCCAGCGAGCATAGCCAAGTTAGGCACCTGCACACATTGCCACTCTTTCTTTTCCTGTCGACG
CTTCAATTTCTTTTACAAACCATCGCAGCGTGTAAT

**nhomie WT**

CTAGCTTGTGGGATGGCCAGGGGGTGTTCTAAGGATGCGTTTATCTGCGCTGAGGATGTGCTAAT
TTTTTACTCAGCAGACCTTTCTGCGAACAGAAATTGAGCACTGAGTGCGCAGGGTGGGGGTGAA
TTGAATCATCTTTGAATGACTTTTGTTCCTTGGGATTTTTTCAATAGATTAGGATGGACTTTTACCATT
CAAAATACAATACTATTAATCTTAAAAGTTACTTATGTTACTTAAGCAAATTAAGCGGTTAAGTGGAGT
CCAAATGGCGGTCAACATTTATATGTAGAAATAAATTCGTAAATTCGTAAATTCGATAAACTTATAAG
GTGTTTTGATTATTTCTAATATTTGTTAGATAAGATGAATAATTTATACTCTATTTTATATACAGTGAACCC
AATTTTTATTTACATATGTGTATGCCCGACTCATGTGGGAAAAATGTAAGTAACGTAGCTCAAACGC
AATCTCAAAAGTATGCAACACTTTCTCAAATTACGTAAGCAGCTCTCGTAGACATACGCTCACATC
AGTGAGCTCCTGTCAGCCAGTAAACATGTCCCTTTATCTGGCAACGCTGCTACCCATTAAAT

**nhomie Δmotif-8**

CTAGCTTGTGGGATGGCCAGGGGGTGTTCTAAGGATGCGTTTATCTGCGCTGAGGATGTGCTAAT
TTTTTACTCAGCAGACCTTTCTGCGAACAGAAATTGAGCACTGAGTGCGCAGGGTGGGGGTGAA
TTGAATCATCTTTGAATGACTTTTGTTCCTTGGGATTTTTTCAATAGATTAGGATGGACTTTTACCATT
CAAAATACAATACTATTAATCTTAAAAGTTACTTATGTTACTTAAGCAAATTAAGCGGTTAAGTGGAGT
CCAAATGGCGGTCAACATTTATATGTAGAAATAAATTCGTAAATTCGTAAATTCGATAAACTTATAAG
GTGTTTTGATTATTTCTAATATTTGTTAGATAAGATGAATAATTTATACTCTATTTTATATACAGTGAACCC
AATTTTTATTTACATATGTGTATGCCCGACTCATGTGGGAAAAATGTAAGTAACGTAGCTCAAACGC
AATCTCAAAAGTATGCAACACTTTCTCAAATTACGTAAGCAGCTCTCGTAGACATACGCTCACATC
AGTGAGCTCCTGTCAGCCAGTAAACATGTCCCTTTATCCGTCGACGCTGCTACCCATTAAAT
