## Supplementary File S2 for "Sequence-to-function modeling uncovers the context-specific grammar of *Drosophila* chromatin insulation": Description of Supplementary File S2.pdf

In Supplementary File S2, we include files used to match Motif-8 and AAG3, two Domino-derived insulator motifs, against DNA-binding motif predictions for four-zinc-finger arrays (4ZF) of C2H2 ZF domains (see .meme file). TOMTOM was used to align the Domino motifs to the 4ZF motif predictions, generating p-values for each match. Benjamini-Hochberg correction was used to generate final q-values. Comprehensive results for all matches are included as separate .tsv files.

Below, each Domino-derived motif (top) and its best corresponding 4ZF PWM match (bottom) are shown. For each 4ZF, the isoform and the particular zinc finger numbers are indicated.

### Motif-8

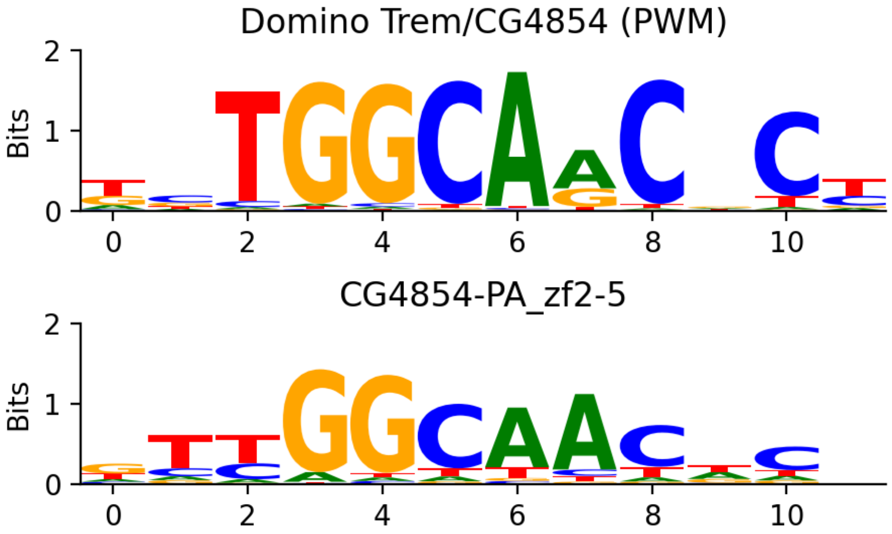


### AAG3

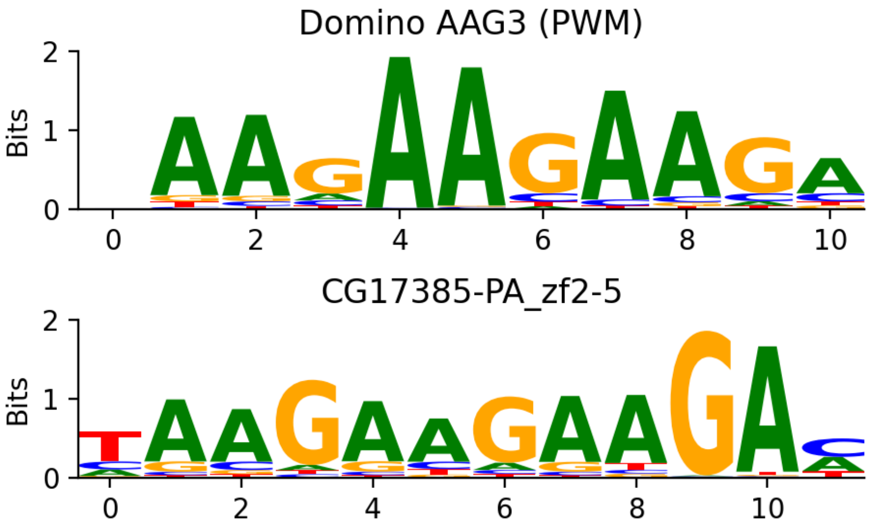
